# Systematic functional annotation of thousands of BAHD acyltransferases in plant genomes using Protein Language Model and phylogenomic tools

**DOI:** 10.64898/2026.06.09.730906

**Authors:** Nathaniel S. S. Smith, Xinyu Yuan, Chesney Melissinos, Sahaj Satani, Crystal Grissom, Gaurav D. Moghe

**Affiliations:** Plant Biology Section, School of Integrative Plant Science, Cornell University, Ithaca, NY, USA

## Abstract

The functional annotation of plant genes lags significantly behind their genomic annotation. Closing this gap requires thorough cataloging of reported protein activities alongside predictive methods that scale beyond sequence-similarity inference. Focusing on the BAHD acyltransferase enzyme family as a model, we assembled FuncZymeDB-BAHD, a large database of 2,705 LLM-retrieved and curated enzyme-acceptor-donor activities covering 336 BAHDs from 156 plant species, a 2-to-6-fold expansion over Swiss-Prot and prior compilations. We further developed FuncPred-OG, which maps queries to orthologous groups and previously characterized enzymes in FuncZymeDB-BAHD, returning hits with high evidence provenance. FuncPred-OG enabled functional prediction of over half of BAHDs across 85 plant proteomes, of which five novel predictions were validated via *in vitro* assays and recent studies. For the remaining BAHDs without FuncPred-OG annotation, we developed FuncPred-AI, where logistic-regression classifiers trained on protein language model embeddings achieved high Area-Under-the-Precision-Recall-curve (AUPR) scores and correct-hit rates up to 93%. FuncPred-AI yielded ≥1 probable donor/acceptor annotation for 99.9% (8894/8897) of BAHDs in our pan-plant dataset. Finally, the FuncPred workflow and datasets were deployed on a web portal for broader utilization, potentially reducing experimentalists’ efforts for selecting candidates from days to minutes. Overall, this framework provides a generalizable template for functional annotation of entire enzyme families.

## INTRODUCTION

Over 4000 plant genomes have been sequenced^1^, each containing tens of thousands of genes. Unfortunately, even in genomes of reference plant species such as *Arabidopsis thaliana*, the functions of less than half of the genes are experimentally determined^2^. This “function gap” represents one of the most significant bottlenecks in translating genomic information into biological knowledge. A central challenge in resolving the function gap is the presence of gene families such as enzyme families. From a basic science perspective, understanding the catalytic repertoire encoded by a genome is essential for reconstructing metabolic networks, elucidating biosynthetic pathways for specialized metabolites, and understanding the molecular basis of plant adaptation^3,4^. From an applied standpoint, knowledge of enzyme function is a prerequisite for metabolic engineering efforts aimed at producing valuable natural products, improving crop quality traits, and developing sustainable agricultural practices^5^. However, most enzymes in plant genomes are poorly or incorrectly annotated with their utilized substrate classes^6^, because of processes such as duplication-divergence, allelic divergence and promiscuity that lead to the failure of function transfer using sequence similarity^7,8^

The BAHD acyltransferase family exemplifies this challenge. Named after the first four biochemically characterized members – BEAT, AHCT, HCBT, and DAT^9^ – this enzyme superfamily catalyzes the transfer of acyl groups from CoA thioesters to a diverse array of acceptor substrates, producing esters and amides that are central to plant defense, signaling, reproduction and development^10,11^. The family expanded dramatically during land plant evolution, from fewer than five copies in algae to ∼100 copies in typical diploid angiosperm genomes^12^. This expansion, driven by gene duplication and sub/neofunctionalization, has enabled plants to produce a remarkable diversity of acylated metabolites including volatile esters, acylated flavonoids and anthocyanins, alkaloid precursors, and cutin and suberin monomers. However, the functional characterization of BAHDs has not kept pace with their genomic proliferation: in a representative crop genome such as tomato, fewer than 15 out of approximately 100 BAHDs have associated substrate class annotations^12^ with questionable or nonexistent provenance.

A major factor underlying the function gap is the “biocuration bottleneck”. Thousands of enzyme activities have been experimentally characterized and reported in the primary literature over the past four decades, yet only a fraction of these have been deposited in structured databases such as UniProt/Swiss-Prot, BRENDA, and KEGG^13^. Our previous analysis revealed that over 70% of experimentally characterized BAHD activities remain uncurated in the public domain^6^. This gap arises partly because biocuration – the labor-intensive process of reading papers, extracting structured information, and validating entries – is performed by a very small workforce, with fewer than 100 full-time equivalent biocurators working across all functional databases and fewer than 10 focusing on bacteria and plants^13^. This bottleneck is further compounded by the exponential growth in published literature, making manual curation of every relevant paper increasingly infeasible.

Multiple computational approaches have been developed to address the enzyme function prediction problem. Traditional methods rely on sequence similarity searches, orthology inference, and phylogenetic analysis to transfer annotations across species, using tools such as BLAST, PAINT, TreeGrafter and PhyloGenes^14–17^. More recently, machine learning and deep learning methods have been applied to predict Gene Ontology (GO) categories, Enzyme Commission (EC) numbers, and enzyme-substrate interactions from protein sequence features^18–20^. Contrastive learning approaches such as CLEAN have demonstrated strong performance in EC number prediction by learning functionally discriminative sequence embeddings^21^. Multimodal approaches that jointly model enzyme and substrate representations have also shown promise; for example, FusionESP integrates protein and chemical language models through a contrastive learning framework to predict enzyme–substrate pairs, achieving state-of-the-art accuracy while demonstrating improved generalization across enzyme families^22^. EZSpecificity is another tool that aims to distinguish closely related enzymes that act on different substrates within the same reaction class^23^ through combinatorial modeling of enzyme and substrate structures. EnzymeCAGE leverages structural and sequence information and graph neural networks to predict whether an enzyme can catalyze a given reaction^24^. These tasks remain particularly challenging for large, functionally diverse families and present challenges for enzyme candidate selection by experimentalists from a long list obtained from RNA-seq or other high-throughput experiments. For example, EnzymeCAGE, depending on the task, reported a 42-58% success in identifying the correct enzyme-reaction pair in its top 10 predictions^24^, and this rate itself may be affected by limited knowledge of promiscuous enzyme reactions. Furthermore, because chemical structures need to be provided as inputs, such approaches may not be suitable for high-throughput functional annotation of enzymes.

Many of these approaches utilize protein language models (PLMs) that generate rich, biologically meaningful representations of proteins. The Evolutionary Scale Modeling (ESM) family^25,26^ and ProtTrans^27^ have been shown to capture functional and structural information directly from amino acid sequences, enabling EC number assignments and substrate chemical structure prediction^28,29^. However, while these methods are very powerful, the quality of all these prediction methods is fundamentally constrained by the breadth and accuracy of the training data available – a limitation that circles back to the curation bottleneck described above. Models trained on incomplete or biased reaction databases risk propagating incorrect annotations, underscoring the need for comprehensive, well-curated enzyme activity datasets as a foundation for reliable computational prediction.

In our previous study^6^, we leveraged recent advances in large language models (LLMs) for accelerating biocuration. We previously developed FuncFetch, a workflow that integrates NCBI E-Utilities, GPT-4, and Zotero to screen thousands of published manuscripts and extract enzyme–substrate information at scale. Applied to nine plant enzyme families, FuncFetch retrieved over 32,000 entries from 5547 papers, with a substrate-level precision/recall of 0.86/0.64 when benchmarked against a manually curated BAHD dataset. While FuncFetch demonstrated the feasibility of LLM-assisted literature mining for biocuration, the extracted data required substantial manual verification before it could be used to train predictive models or populate public databases. The convergence of this large-scale literature mining capability with advances in PLMs presents an unprecedented opportunity to comprehensively annotate an entire enzyme family at scale.

In this study, we leverage this convergence to construct a comprehensive functional annotation framework for the BAHD acyltransferase family across the plant kingdom. Starting from FuncFetch-extracted data, we performed extensive manual curation to assemble FuncZymeDB, a database of 2705 well-curated BAHD–substrate activities – representing a 2-6X expansion over currently available activities in Swiss-Prot and other public databases. In the FuncPred-OG framework, we then exploited this taxonomically rich dataset and phylogenetic relationships among characterized and uncharacterized BAHDs to propagate substrate class annotations, increasing the proportion of BAHDs with predicted activities from <10% to ∼50% across 85 plant species. For the remaining uncharacterized enzymes, we developed FuncPred-AI. Specifically, we trained classifier models to predict acceptor and donor substrate categories directly from protein-derived embeddings from protein language models (PLMs), achieving high accuracy. Finally, we integrated these annotations into a publicly accessible web resource encompassing 8897 BAHDs from 85 plant species, complete with a user-facing tool for predicting functions of novel BAHD sequences. Together, this work establishes a generalizable paradigm – from literature mining (FuncFetch) to curated database (FuncZymeDB) to phylogenomic inference (FuncPred-OG) to deep learning-based prediction (FuncPred-AI) – for dramatically closing the enzyme function annotation gap in plants. Importantly, this framework allows experimentalists to rapidly select candidate BAHDs for further biochemical studies, potentially reducing the timescale for such selection process from days and weeks to minutes.

## RESULTS

### The BAHD activity space is very diverse

We first manually curated and analyzed the diversity of entities and enzyme-substrate relations in the extracted FuncFetch BAHD dataset, and – along with additional manually reviewed activities – added them to our previously published gold standard set^6^. For curation, every paper was read carefully by at least two co-authors to verify extracted entities. Additional attributes such as SMILES, InChiKey, CHEBI-ID, and NPClassifier categories^30^ were connected to the substrates. This fully curated dataset **(Supp. File 1A)** contained 2705 individual entries, each entry corresponding to a specific enzyme-acceptor-donor relationship from a given paper and a given species **(Supp. File 1A)**. This BAHD dataset, which we refer to henceforth as FuncZymeDB-BAHD or FZ-BAHD, represents the reaction space of all BAHDs gathered manually or extracted by FuncFetch and formed the basis of all our downstream analyses. FZ-BAHD contains 431 reported BAHDs corresponding to 336 unique sequences from 156 species (**Fig. 1A**), utilizing 384 unique acceptors and 73 unique donors (**Fig. 1B, Supp. File 1B)**. These, together, represent 2247 unique enzyme-acceptor-donor reactions. Compared to BAHDs in Swiss-Prot and our previous publication^12^, our BAHD dataset contains 2-6-fold additional entities (**Table 1**).

**Figure 1:**
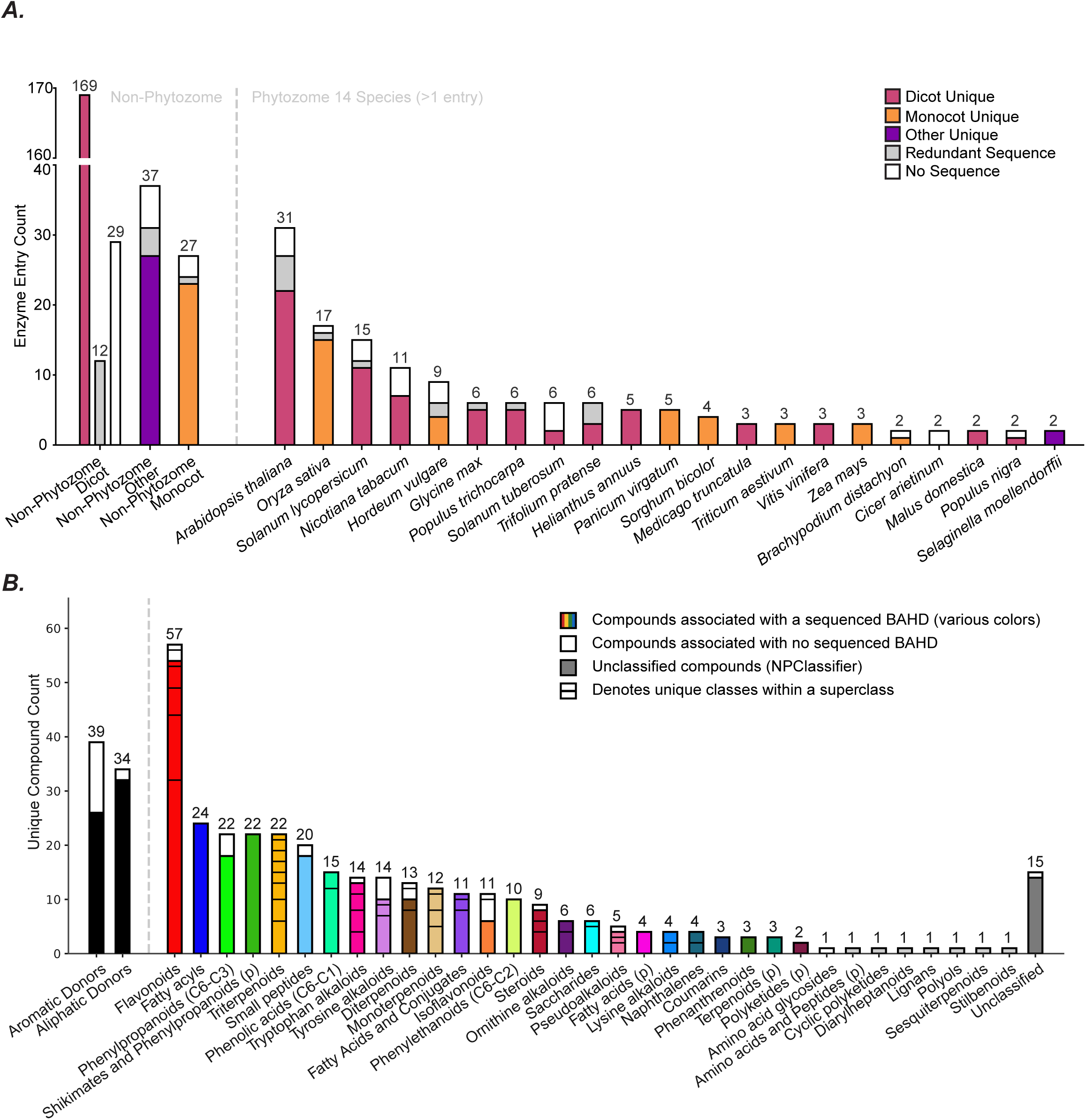
BAHD Sequence & Substrate Summary. **(A)** BAHD Entries by Species and Sequence Status. List of species extracted from Phytozome 14 website and cross-referenced against FuncZymeDB. Redundant sequences are identical proteins from the same species and are removed from the total for the count of unique sequences. Note the y-axis break and non-stacked bars for Non-Phytozome Dicot entries. **(B)** Counts of BAHD substrate donor types and acceptor NPClassifier categorizations. Unique compounds determined by SMILES. Acceptor categories are pathways where noted by (p), otherwise superclasses. Horozontal bar breaks denote seperate NPC classes within each superclass. Individual compounds may have more than one class/superclass, and are counted in each independently.

**Table 1:** Count of unique entities available in various sources, as of Nov 2025.

| Entity | # in Swiss-Prot 2025_11 | # in Kruse et al, 2022 | This study | Fold change vs. Swiss-Prot 2025_11 | Fold change vs. Kruse et al, 2022 |
| --- | --- | --- | --- | --- | --- |
| Papers | 64 | 106 | 213 | 3.3X | 2.0X |
| Species | 36 | 74 | 156 | 4.3X | 2.1X |
| **BAHDs with substrate information** | 70 | 164 | 336 | 4.8X | 2.0X |
| **Acceptors** | 62 | 206 | 384 | 6.2X | 1.9X |
| **Donors** | 25 | 31 | 73 | 2.9X | 2.4X |

Categorization of substrates into broad groups can simplify inferences and align with biological reality of substrate promiscuity of BAHDs^12^. To accomplish this, BAHD substrates in FZ-BAHD were sorted into 71 Classes, 29 Superclasses, and 7 Pathways using NPClassifier (NPC) (**Fig. 1B**)^30^. Thirty-two unique acceptors (8.3%) were assigned pathway level annotations alone, and 14 (3.6%) were unassigned. When a Superclass was unavailable, Pathway annotations were used as a fallback annotation throughout this work. BAHDs were found to be associated with every Pathway and 37.7% (29/77) of Superclasses in the NPC ontology, demonstrating the large substrate space occupied by this family^30^. All donors were assigned to the same NPC Class Fatty acyl CoAs **(Supp. File 1B)**, but BAHDs can have preferences towards aliphatic or aromatic CoAs^31–33^. Therefore, we categorized donors into two types: aliphatic or aromatic, containing 34 and 39 unique donors, respectively (**Fig. 1B**). On the acceptor substrate side, most of the extracted reactions use flavonoids (involved in stress response and mutualistic interactions), followed by various fatty acyls (e.g. long-chain acyl-CoAs; involved in cuticle biosynthesis and pathogen defense), shikimates and phenylpropanoids (e.g. shikimate; lignin biosynthesis), small peptides (e.g. spermidine; stress response) and triterpenoids (e.g. oleanolic acid; stress response). (**Fig. 1B; Supp. File 1B)**. Combinatorial acceptor-donor analysis (**Fig. 2A**) clearly illustrated the ubiquitousness of acetyl-CoA as a donor in several BAHD-mediated reactions, primarily for mono-, di- and tri-terpenoids, fatty acyls and flavonoids. Physiologically, these acetylated compounds tend to be in pathways important in reproduction and stress response^34–36^. Phenylpropanoids and different alkaloids tend to share multiple aromatic CoAs as donors, while malonyl-CoA is preferred for Flavonoid acceptors. Compared to these donors, use of other aliphatic CoAs in biochemical studies is low and acceptor-specific. To visualize the relationship between substrates, we constructed a gravity-weighted network of observed acceptor-donor relationships. Edges were weighted by the number of unique BAHD-catalyzed reactions between specific donor types and acceptor categories (**Fig. 2B**) This network revealed that the five highest weighted connection groupings were between malonyl-CoA and flavonoids (43 reactions), acetyl-CoA with triterpenoids and fatty acyls (21 and 20 reactions, respectively), and caffeoyl/p-coumaroyl-CoA and Flavonoids (20 and 19 reactions, respectively) **(Supp. File 1C)**. A large number of nodes at the network edges also reinforced the diversity of unique acceptor-donor reaction combinations, highlighting the promiscuity and divergence of BAHDs.

**Figure 2:**
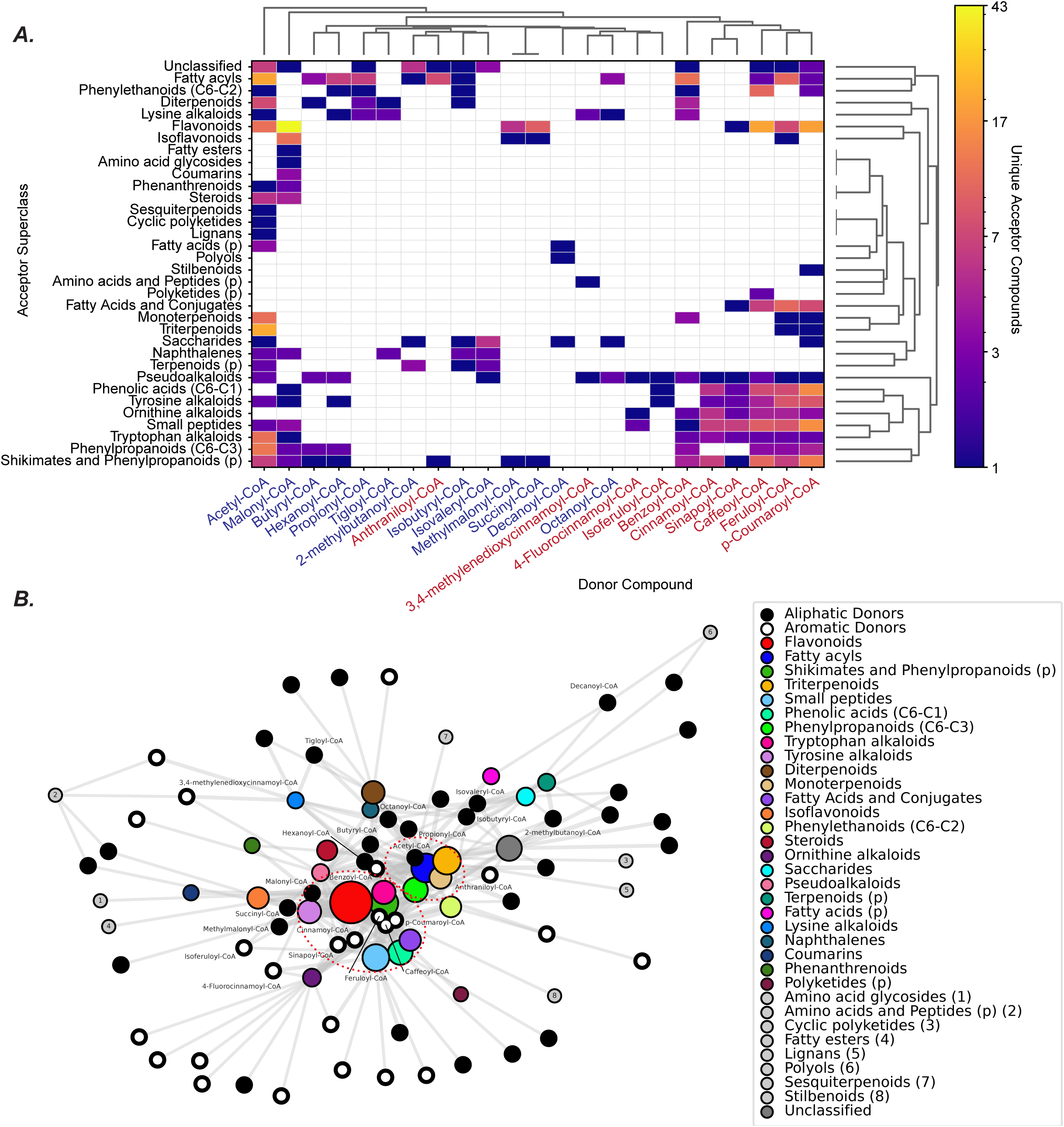
BAHD Reaction Space. **(A)** BAHD reaction heatmap. A donor-acceptor superclass matrix was generated from unique acceptor compound counts for each donor-superclass pair. Rows and columns were independently clustered using SciPy average-linkage hierarchical clustering. Counts were logarithmically normalized before color mapping, with zero counts shown in white. To reduce visual complexity, only donors connected to at least three unique acceptor superclasses are shown; same donor set is labeled in 2B. Aliphatic CoA donors are labeled in blue and aromatic CoA donors in red. **(B)** BAHD reaction network. Donor nodes represent individual donor compounds, and acceptor nodes represent acceptor superclasses or pathway fallbacks (denoted with “(p)”). Edges indicate observed donor-superclass relationships, with edge width and layout attraction scaled linearly to the number of unique acceptor compounds included in each pair. Network drawn using NetworkX spring_layout with the energy parameter set. Acceptor node area was scaled by the number of unique acceptor compounds in each superclass. Acceptor nodes with a single unique compound are numbered and shown in light grey for display. Red dashed ovals highlight two major reaction-pair clusters.

FZ-BAHD contains enzymes from 156 different plant species, with *Arabidopsis thaliana*, *Oryza sativa* (rice) and *Solanum lycopersicum* (tomato) being the largest contributors. Most of the sequenced BAHDs (65.2%, 219/336) are from 111 species without a reference genome in Phytozome 14 (**Fig. 1A**). A long tail of 79 species has only a single characterized BAHD **(Supp. File 2)**. These dataset characteristics highlight the research community’s interest in investigating the lineage-specific variation in plant metabolism.

### Mapping of BAHDs to orthologous groups enables substrate class prediction for over half of the BAHD enzyme space

The high diversity of species, enzymes and substrates in FZ-BAHD creates an opportunity to predict functions of uncharacterized BAHDs using sequence similarity. First, through phylogenetic reconstruction, we confirmed the presence of previously defined BAHD clades in our dataset (**Fig. 3**)^10–12,37^. Clade 2, which comprised genetically characterized enzymes CER2 and Glossy2 at the time of data analysis, was not represented in this study. The acceptor substrate distributions in each of the other clades were consistent with our previous analysis^12^. Two clades contain BAHDs that broadly condense aromatic donors with phenolic acid (C6-C1) (Clade 5: 71%, involved mostly in lignin biosynthesis) or ornithine alkaloids and pseudoalkaloid acceptors (Clade 4: 89%/68%, respectively, involved mostly in polyamine and alkaloid metabolism). Another two clades were observed mostly donating aliphatic groups to flavonoids or shikimates and phenylpropanoids (C6-C3) (Clade 1: 65% [flavonoid biosynthesis], Clade 7, 100% [coniferyl/cinnamyl alcohol acylation], respectively). Clades 3 and 6 interact with a greater mix of acceptor categories and donor types, however, at the sub-clade levels, substrate class clustering was seen among related enzymes (**Fig. 3**).

**Figure 3:**
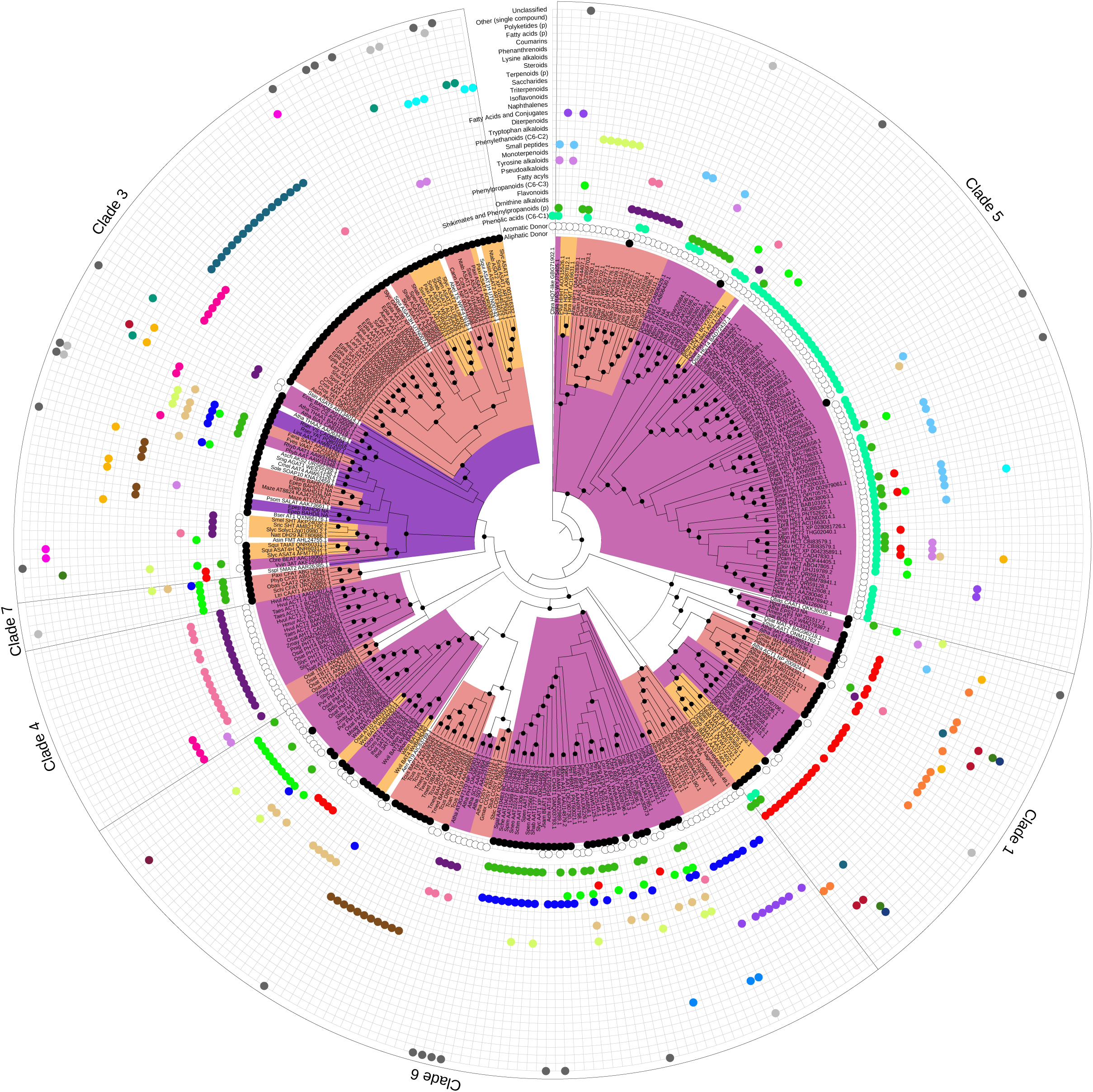
BAHD Phylogeny and Substrate Annotations. Colored ranges on the branches indicate individual OGs from their last common ancestor to the extant enzymes in those OGs. Labels with white background are BAHDs in unique, separate OGs. Tree was generated using IQ-TREE3 (1000 UF bootstrap replicates). Black circles on tree nodes indicate UF bootstrap values >90. Each concentric circle represents a ontological category obtained using NPClassifier.

FZ-BAHD can be used to predict the functions of novel, uncharacterized BAHDs, however, many plant clades are unrepresented in FZ-BAHD species. To enhance the predictability of substrate classes based on phylogenetic relationships, we combined FZ-BAHD with BAHD sequences from high-quality proteomes of 85 plant species **(Supp. File 4)** and used them to establish orthologous groups (OGs). In our previous study^12^ – by validating OG-derived predictions of 34 novel BAHDs via experimental and literature analyses, we estimated an 80-90% success rate in identifying the correct substrate class of novel enzymes. We further used the OGs to define the “narrowest common lineage” and the “lineage levels” that identify the depth of conservation of each OG (**Fig. 4A**). For example, an OG with BAHDs from only Solanaceae-family species will have the narrowest common lineage as “Solanaceae” and the lineage-level as “Family” (**Fig. 4B**). We note that these two metrics, while useful for interpretation and enzyme prioritization, are limited by the species representation in our OGs; as we include more species, species-level OGs could transition to genus or family-level OGs, therefore interpretation of these two metrics should be made carefully. Overall, the 336 BAHDs were assigned to 54 OGs using BLAST. Across all OGs, 184 have their lineage-level beyond “Order” indicating their conservation at broad phylogenetic scales. However, 148 (80.4%) of these OGs do not have a single characterized enzyme activity (**Fig. 4A**). For example, there are 5 BAHDs in *Solanum lycopersicum* that map to OGs with “Tracheophyte” as the narrowest lineage (**Fig. 4B**). This finding highlights the many conserved and likely important BAHDs that remain uncharacterized.

**Figure 4:**
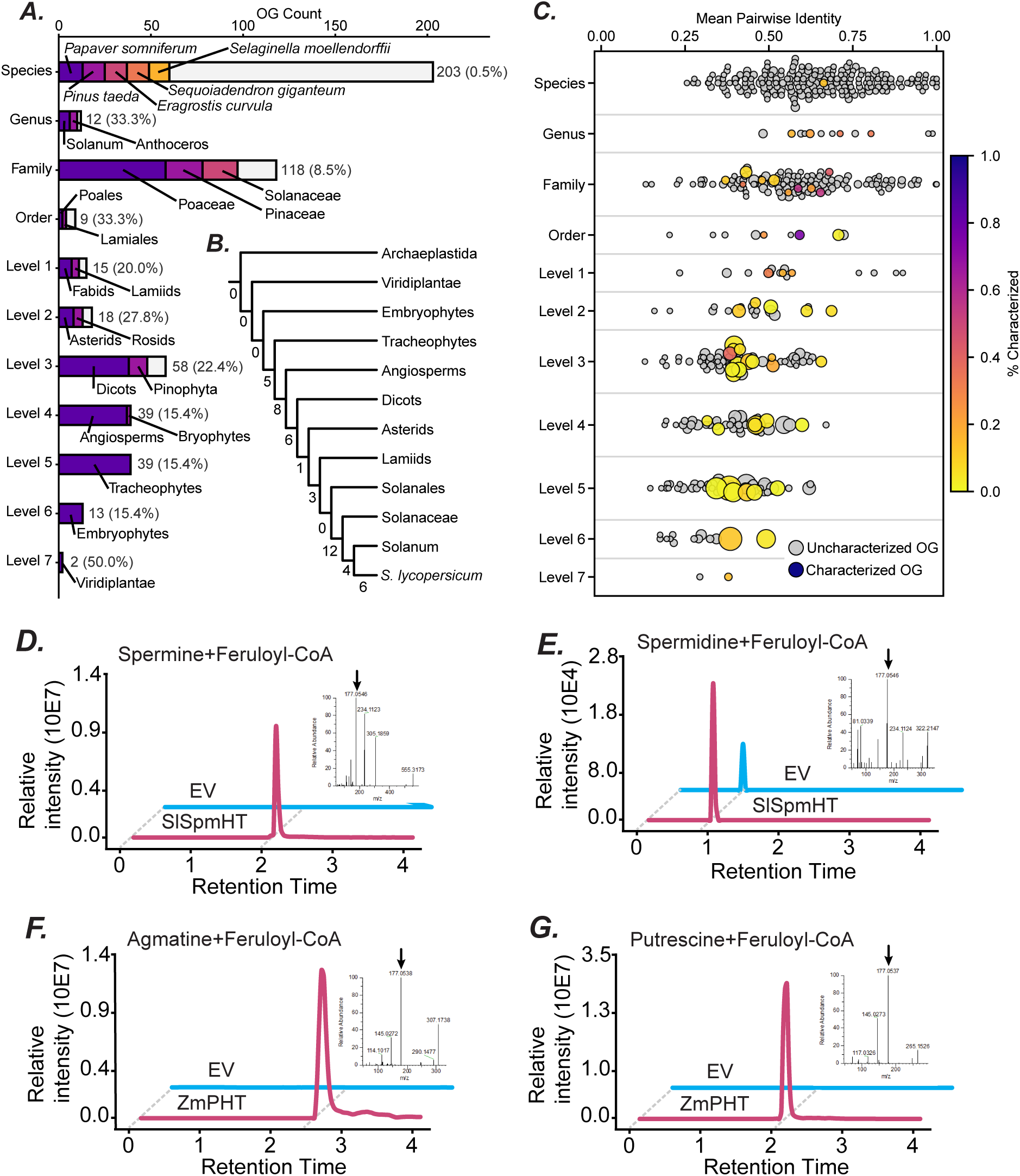
BAHD Orthologous Groups (OGs). **(A)** OG Counts by Narrowest Lineage Level. The most specific common lineage of all BAHD sequences within an OG was determined and logged in the OG database module (”narrowest_common_lineage”). Clade levels broader than Order were defined manually and range from Class to Kingdom. The top 1-5 lineages by count per clade level are highlighted and labeled. Total OG count and percentage of OGs at a given level with at least one characterized BAHD are noted to the right of the bars. **(B)** Simplified Cladogram of *S. lycopersicum* BAHDs. Node labels are counts *S. lycopersicum* sequences in uncharacterized OGs with narrowest common lineage of the upper branch. **(C)** OG Mean Pairwise Identity by Narrowest Lineage Level. Pairwise Identity calculated as fraction of aligned residues with exact non-gap matches (biopython Bio.Align.PairwiseAligner.align). Circle area scaled linearly with count of sequences in OG. **(D)** Chromatogram of SlSpmHT Product Diferuloyl Spermine. Quantifier ion m/z: 177.0546. **(E)** Chromatogram of SlSpmHT Product Feruloyl Spermidine. Quantifier ion m/z: 177.0546. **(F)** Chromatogram of ZmPHT Product Feruloyl Agmatine. Quantifier ion m/z: 177.0538. **(G)** Chromatogram of ZmPHT Product Feruloyl Putrescine. Quantifier ion m/z: 177.0537.

Among BAHD clades, greater sequence similarity is associated with greater substrate class similarity (R^2^=0.81, t-test p=0.0136; **Supp. File 5A**). The same association, albeit weaker, was observed across OGs, indicating that sequence and substrate similarity covary at broad clade and fine OG scales (R^2^=0.22, t-test p=0.0048; **Supp. File 5A**). Furthermore, OGs with narrower lineage levels tended to have greater sequence similarity, reflecting more recent divergence (**Fig. 4C**). These results also revealed the large number of OGs without any experimental evidence. We expect that such a bias will be present for other enzyme families too.

Candidate enzyme prioritization for pathway discovery and genotype-to-phenotype mapping frequently involves phylogenetic reconstruction and comparisons with activities of previously characterized homologs – a time and labor-intensive process. To expedite this step for BAHDs, we designed a web tool (FuncPred-OG; https://tools.moghelab.org/funczymedb) which automates mapping of novel BAHD sequences to OGs. FuncPred-OG then provides an output of any characterized enzymes in that OG as a ranked list, along with known substrates, narrowest lineage and high-resolution evidence provenance. We processed 8897 BAHD sequences from 85 plant proteomes with FuncPred-OG, enabling mapping of 4743/8897 (53.3%) of all BAHDs to OGs with characterized enzyme activities. To further test the accuracy of these OG-based predictions beyond our previous study^12^, we cloned and functionally characterized BAHDs from tomato (*Solanum lycopersicum*), maize (*Zea mays*), and rice *(Oryza sativa)* **(Supp. File 6A, B).** Solyc12g010980 lies in the same OG as its best BLAST hit *Solanum richardii* SpmHT (SrSpmHT; AMB21755.1) at 70.6% identity **(Supp. File 6C)**. SrSpmHT is annotated as spermine hydroxycinnamoyl transferase using spermine and feruloyl-CoA as substrates^38^. Solyc12g010980 also showed the ability to utilize these substrates *in vitro* (**Fig. 4D**) and therefore was annotated as SlSpmHT. SlSpmHT also exhibited weak activity toward spermidine as acyl acceptor, indicating a broader substrate utilization (**Fig. 4E**). Using FuncPred-OG predictions, we also characterized *Zea mays* enzyme Zmay_GRMZM2G095327 **(Supp. File 6E)** and annotated it as a putrescine hydroxycinnamoyl transferase (ZmPHT), based on its reaction of feruloyl-CoA with agmatine and putrescine to produce feruloyl agmatine and feruloyl putrescine, respectively (**Fig. 4F, G**). Finally, two additional enzymes could not be experimentally validated in our *in vitro* enzyme assays; however, other groups published their activities recently, and these are consistent with our predictions **(Supp. File 6D, F)**. First, the tomato protein Solyc02g093180 maps to an OG containing four known enzymes Atha_EPS1, Amem_AT7-3, Lsat_BOS, and Hpol_BAHD3, and its activity with isochorismoyl-L-glutamate – a substrate used by Atha_EPS1 – was experimentally validated recently^39^. Second, the rice enzyme Os10G35950 was also shown recently to use benzyl alcohol and benzoyl-CoA as substrates^40^, consistent with its mapping to an OG with similar activities, leading us to annotate it as benzoyl-CoA:benzyl alcohol benzoyltransferase (OsBEBT).

This experimental validation builds on the 34 activities we used for validation in our previous study^12^ and further confirmed the feasibility of using FuncPred-OG for identifying candidate substrates for experimental testing. However, FuncPred-OG was able to predict functions for only 53.3% of the analyzed plant BAHDs. The remaining 46.7% of BAHDs map to OGs without any characterized enzyme and hence, their substrate classes cannot be predicted. Therefore, we developed FuncPred-AI, an approach that uses protein language models (PLMs) for substrate class prediction without a need for knowledge from homologous proteins.

### Protein language model representations provide signal of BAHD function

We evaluated the performance of two state-of-the-art PLMs, Evolutionary Scale Model Cambrian (ESMC) & ESM3^20,41^, on the FZ-BAHD dataset. ESMC uses only sequence as input whereas ESM3 is multimodal, accepting sequence, structure and function tracks. However, the standard output of these features is InterPro annotations, which is not informative for predicting enzyme substrate classes. Therefore, using our highly curated BAHD dataset as gold standard, we developed logistic regression classifiers to classify enzymes as using acceptor and donor classes, using feature sets derived from the PLMs. We compared four model-derived feature sets as inputs to the classifier, namely embeddings derived from only sequence (ESMC:SeqEmbed), sequence and structure (ESM3:StrucEmbed), function logits (ESM3:FuncLogit), and a concatenation of the function logits with the structure embedding (ESM3:FL-SE).

As an initial test of whether each feature set contained functionally relevant variation, we performed Principal Components Analysis (PCA) on the raw values from each feature set (without any training/classifier) and quantified separation by OG and donor type via Fisher discriminant ratio. We included OG because it displayed functional signal earlier in phylogeny analysis, and donor type because it is a label shared across much of our dataset. We found that all four inputs carried information to distinguish BAHDs (**Fig. 5A, Supp. Fig. 1, 2).** Notably, the ESM3 function logits provided greater discrimination of OGs and donor types than either the sequence or structure embeddings (**Fig. 5A, Supp. Fig. 1, 2)**, with PC1 explaining >99% of the variation in the BAHD dataset **(Supp. Fig. 1, 2)**. However, this signal was reduced when inputs were regularized, suggesting that the magnitude of feature variation contributed to the observed separation (**Fig. 5A**). Nonetheless, we subsequently trained substrate-specificity classifiers using all four feature sets as inputs to directly test the transferability of these representations to our BAHD substrate prediction task.

**Figure 5:**
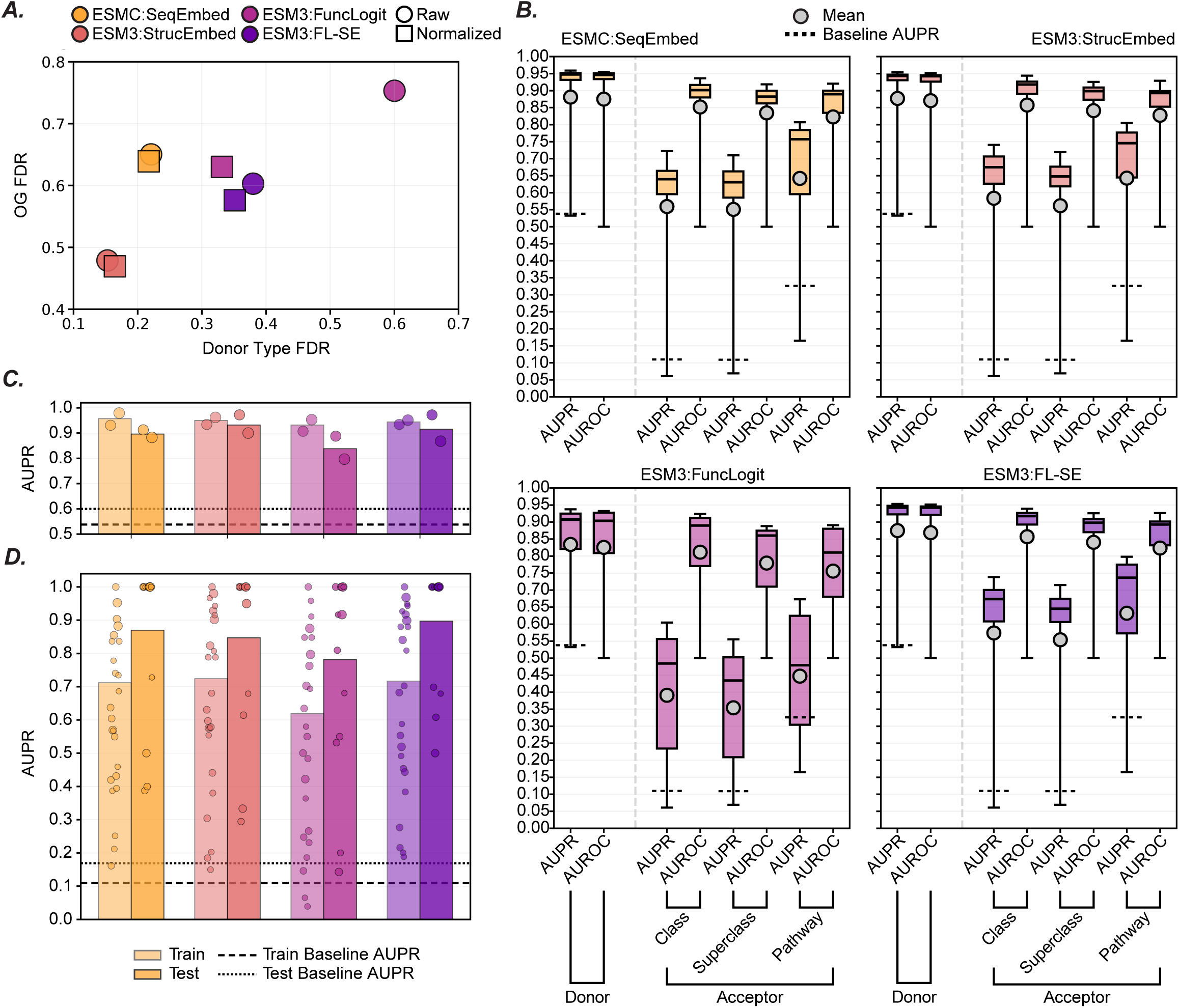
Optimization of FuncPred-AI. **(A)** Fisher discriminant ratio (FDR) of OG and donor type by input feature. One-vs-rest FDR was calculated in the full-dimensional principal component space and summarized as the unweighted mean across categories. L2 normalization was applied to raw features where indicated. **(B)** Logistic regression classifier parameter optimization. Solver, regularization penalty, inverse regularization strength (C), and tolerance were optimized using 5-fold GridSearchCV.The best-performing parameters were frozen and used for all subsequent training runs. Baseline AUPR was calculated by evaluating deterministic sklearn DummyClassifier strategies (most_frequent, prior, constant_0, constant_1) for each one-vs-rest label, selecting the best-performing strategy by AUPR, and aggregating across labels using the same label-weighting scheme as the corresponding model metric. The same baseline procedure was applied in 5C and 5D. **(C)** Donor classification performance. Label-size-weighted AUPR is shown for training and test sets. Individual one-vs-rest label AUPRs are overlaid as circles scaled by the number of positive training examples for each label. Logistic regression classifiers were trained using the frozen hyperparameters from 5B. Training performance was estimated by leave-one-out cross-validation, and test performance was evaluated using final models fit on the full training set. Only labels with at least one positive example in the test set were included in test-set AUPR aggregation. **(D)** Acceptor class classification performance. Same as 5C, but for acceptor class labels.

We trained logistic regression classifiers to predict broader labels: donor type (aliphatic/aromatic) and acceptor NP class of BAHDs (**Fig. 1B**). After parameter tuning across 5 cross-folds, all feature set inputs achieved Area Under the Precision Recall curve (AUPR) values exceeding 0.9 on donor type prediction, with ESMC:SeqEmbed reaching a maximum value of 0.96 (**Fig. 5B**), significantly greater than the baseline (0.54). For the more difficult acceptor class task, ESM3:StrucEmbed and ESM3:FL-SE yielded the highest AUPR values of 0.74 and 0.74, respectively, each outperforming the baseline (0.11) substantially. ESM3:FuncLogit reached the lowest maximum AUPR across both tasks and had the greatest range of performance across all tested parameters.

To test the utility of these candidate models, we implemented a leave-one-out cross-validation strategy using fixed per-model parameters that maximized acceptor class AUPR on 5 cross-folds (**Fig. 5B**). This regime was chosen to simulate prediction of a single novel BAHD based on all prior information, and to account for the biological reality of BAHD promiscuity. Most feature sets maintained similar results across training set metrics, apart from ESM3:FuncLogit which scored below 0.7 AUPR on the acceptor class task (**Fig. 5C**). AUPR declined as sequence identity thresholds became more stringent and as the per-label training data decreased **(Supp. Fig. 3A-C)**, a pattern shared with sequence-based methods: both approaches perform best when evaluated cases closely resemble previously characterized examples.

Given the comparable performance on the acceptor class task, we selected ESMC:SeqEmbed and ESM3:FL-SE as candidate classifiers for large-scale BAHD annotation. Analysis of the confusion matrix revealed consistent misclassifications between certain classes **(Supp. Fig. 4A)**. For example, BAHDs with “Branched fatty acids” as acceptors are frequently predicted to utilize “Unsaturated fatty acids” and vice-versa. These errors likely stem from factors including overlapping labels from multi-label compounds and inherent structural similarities between certain classes. To mitigate this ambiguity, we merged classes which exhibited a threshold of mutual activations (**Fig. 6A).** With these refined classifiers established, we proceeded to validate their capabilities on a practical downstream task.

**Figure 6:**
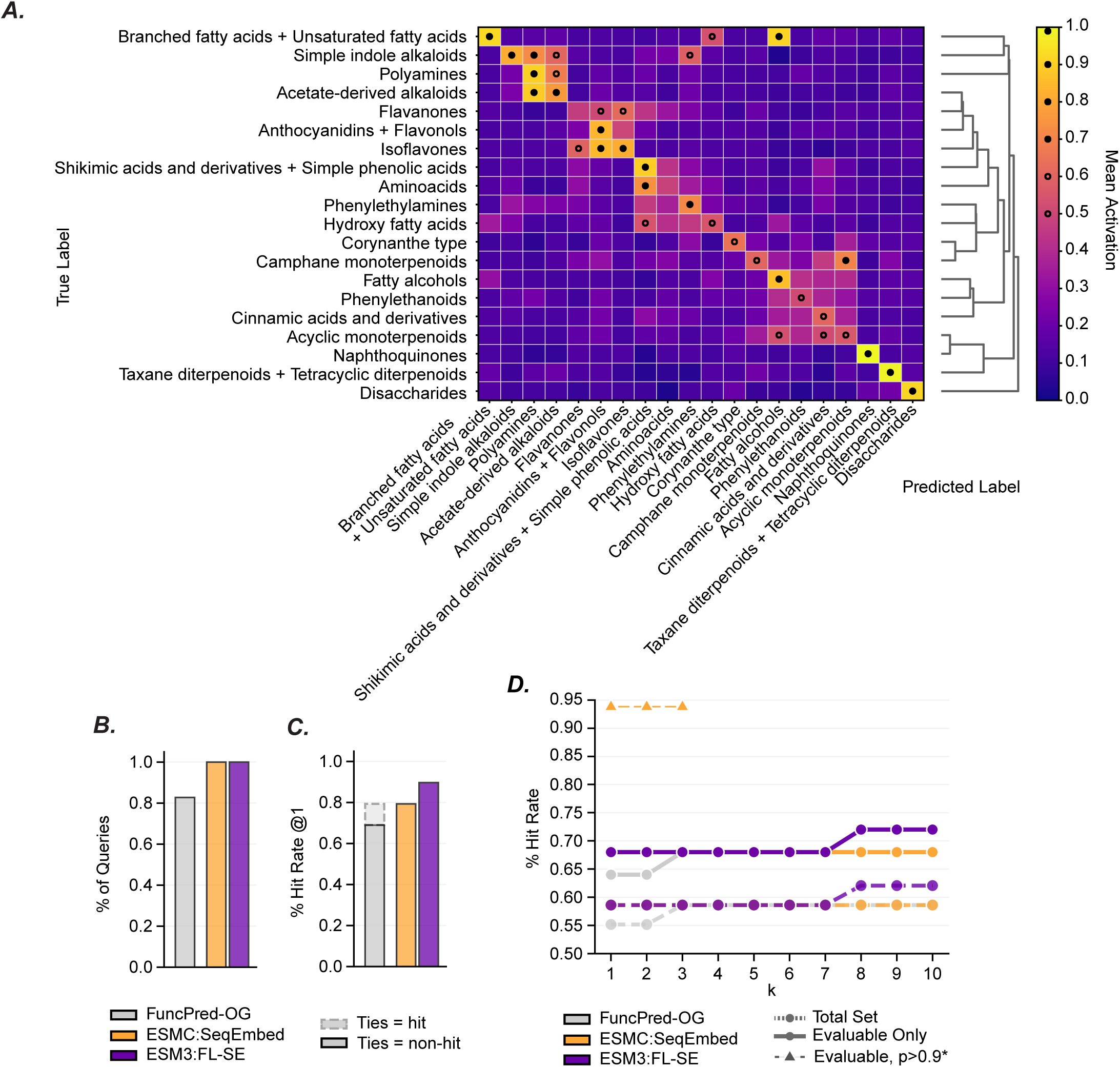
Comparison of FuncPred-AI & FuncPred-OG Figure 6: Comparison of FuncPred-AI & FuncPred-OG. **(A)** Post-correction test-set ESMC:SeqEmbed acceptor class confusion heatmap. Each cell shows the mean prediction activation for the column label among test-set enzymes with the row label. Hollow black circles mark cells with mean activation > 0.5. Filled black circles mark cells with mean activation > 0.7. Acceptor class labels were merged from the pre-correction test-set heatmap in Supp. Fig. 4A when both off-diagonal directional activations for a label pair were >= 0.7. Merged-label prediction scores were collapsed by taking the maximum score among the corresponding pre-merge labels. **(B)** Query coverage across methods. Bars show mean per-task coverage, computed as the fraction of shared comparison queries with a non-empty top-1 prediction, averaged across donor and acceptor class tasks. **(C)** Donor type prediction Hit Rate @1 across methods. The dark bar gives strict Hit Rate @1, where a hit requires a correct unique highest-scoring donor label. The lighter extension gives lenient Hit Rate @1, where ties at the top score are counted as a hit if any top-scoring donor label matches the true donor type. **(D)** Post-correction acceptor class prediction Hit Rate @k on the held-out test set. A hit was defined as at least one true label within the top k predicted labels. Ties at the kth boundary were retained. Solid lines indicate the evaluable-only subset, defined as queries for which at least one true label was present in that method’s possible outputs, and dashed lines indicate the full common query set. The thin, dashed line with top3 points represents the proportion of evaluable set, filtered by acceptor class prediction probabilities of >0.9. Only 16 enzymes form this filtered evaluable set. Acceptor class labels were remapped using the merge groups derived from Supp. Fig. 4A.

### FuncPred-AI improves and extends upon FuncPred-OG

To further augment our test set, we incorporated SlSpmHT and ZmPHT into a held-out set of 27 BAHDs with previously published activities **(Supp. File 7A)**. This expanded collection is characterized by inherent class imbalance; for example, 34.5% (10/29) of the test set enzymes utilize tetracyclic diterpenoids **(Supp. File 7B)**. Because FuncPred-OG and our classifier strategies yield fundamentally different output formats, we compared their respective abilities to rank donor and acceptor labels. Specifically, FuncPred-OG produces sequence-identity-based rankings of validated substrates conversions, which we converted to known labels, whereas our classifier rankings are based on model activation scores for each predicted label.

As expected, FuncPred-OG was limited to predicting only 82.7% (24/29) of the test enzymes, as it was restricted to those with characterized within-OG hits (**Fig. 6B, Supp. File 7A)**. In contrast, both ESMC:SeqEmbed and ESM3:FL-SE achieved 100% coverage across the entire test set, demonstrating the significant expansion in annotation potential offered by PLM-based classifiers. On the donor type task, ESM3:FL-SE achieved the highest accuracy, with 89.7% (26/29) top-1 correct predictions. Accounting for overlapping labels, the hit rate for both FuncPred-OG and ESMC:SeqEmbed was 79.3% (23/29) (**Fig. 6C**). For the more challenging acceptor class prediction task, both classifiers modestly outperformed FuncPred-OG at *k=1*, with ESMC:SeqEmbed and ESM3:FL-SE both reaching a 58.6% (17/29) hit rate compared to 51.7% (15/29) for FuncPred-OG. At higher *k* thresholds, FuncPred-OG’s performance converged with the classifiers at *k=3* (17/29), while the ESM3:FL-SE hit rate rose to 62.1% (18/29) at *k=8* (**Fig. 6D**). These results indicate that the accuracy of the classifier-based and orthology-based approaches are comparable, with donor class predictions of both FuncPred-OG and FuncPred-AI being more accurate than acceptor class predictions. However, FuncPred-AI outperforms FuncPred-OG in scale.

Our final step was to select a candidate classifier for high-throughput downstream annotation. Given the similarity of performance on our validation tasks, we considered the practical trade-offs between predictive performance and computational scalability. While ESM3:FL-SE demonstrated adequate performance on training and validation tasks, its reliance on structural information presented a bottleneck. Feature generation for ESM3:FL-SE on our hardware was consistently more computationally expensive than the sequence-based embeddings of ESMC **(Supp. Fig. 5)**. Consequently, due to the superior scalability of sequence-based inputs and the reduced computational overhead, we selected the ESMC:SeqEmbed-derived classifier as our final model for FuncPred-AI.

Annotation of thousands of BAHDs using FuncPred-OG and FuncPred-AI produced near-universal annotations of 8897 uncharacterized BAHDs **(Supp. Fig. 7)**, representing a massive expansion over the orthology-limited scope of FuncPred-OG. Using a probability threshold of 0.5, FuncPred-AI provided at least one donor or acceptor prediction for 99.9% (8894/8897) of our uncharacterized sequences, demonstrating effectively complete coverage **(Supp. Fig. 6)**. These predictions are available for download on our lab website (https://tools.moghelab.org/funczymedb). Acceptor class assignments were the most comprehensive, covering 8809 sequences (99.0%), while donor type predictions covered 7704 sequences (86.6%). Notably, 7619 sequences (85.6%) received both donor and acceptor annotations.

We further manually verified FuncPred-AI activity predictions of eight BAHDs present in FZ-BAHD and two additional novel BAHDs^42,43^ to check whether they were in the structural neighborhood of the known activities **(Supp. File 7C)**. The donor and acceptor class predictions matched in 10/10 and 9/10 cases. For example, the tomato Solyc12g006330 is annotated as acylsugar acyltransferase^44^ and uses sucrose as the acceptor substrate. FuncPred-AI predicted disaccharides as the substrate class of this enzyme. A newly characterized spermidine caffeoyltransferase from *Lycium barbarum*^43^, which was not in the training data, was predicted correctly to use polyamines and aromatic CoAs. FuncPred-AI correctly predicted the *Neoblechnum brasiliense* HCT5^42^ to use aromatic CoA donors but the probabilities for the top acceptor class predictions were low (∼0.74) compared to >0.95 for the 9/10 correct examples. Reanalyzing the results with the 29 test set enzymes further confirmed that filtering FuncPred-AI predictions using a probability threshold of 0.9 – which reduced the number of enzymes with predictions from 29 to 16 – nonetheless increased the top3 hit-rate from 58.6% to 93.8% (**Fig. 6D, Supp. File 7A)**. These results suggest that high probability FuncPred-AI predictions can effectively narrow down the substrate classes that an experimentalist must test.

Whereas FuncPred-OG was constrained by the availability of characterized orthologous groups, FuncPred-AI extended specificity predictions to nearly the entire dataset, leaving only three sequences without any AI prediction and only one sequence without an annotation from either method. On average, ∼53% of the BAHDs in each species could be annotated using the FuncPred-OG approach with their acceptor and donor classes, while a combination of both FuncPred-OG and FuncPred-AI annotated almost every BAHD in every species (**Fig. 6D, Supp. Fig. 7)**. Both FuncPred-OG and FuncPred-AI are accessible via web-based applications, which allows for the functional inference of novel BAHD sequences provided as user inputs. Together, these resources provide both a systematic annotation resource and an extensible framework for the ongoing functional characterization of BAHD acyltransferases.

## DISCUSSION

This study presents a systematic approach for structuring published information on enzyme activities and applying it towards whole-family functional annotation. We augmented sequence-similarity based function transfer with a PLM-based framework capable of proteome-scale inference and applied this workflow on the BAHD acyltransferase family. While we used BAHDs as a representative large enzyme family for this research, the FuncFetch-FuncZymeDB-FuncPred workflow can serve as an effective template for other enzyme families as well. Our predictive techniques leverage the thousands of characterized enzyme activities published across decades of biochemical research. Previously, we found that most activities retrieved by the FuncFetch workflow were undocumented in the public domain^6^ (**Table 1**). For example, *Piper nigrum* piperamide synthase (PnPAS, P0DO57) has one annotated acceptor-donor pair in Swiss-Prot, compared to 10 unique pairs in FZ-BAHD. Such biocuration gap exists for every enzyme family. To alleviate this challenge, minimally curated FuncFetch data for other enzyme families is available on our website (https://tools.moghelab.org/funczymedb) for download by the research community and can supplement retrieved functional data from other LLM-based approaches^45,46^. We expect that as LLMs continue to rapidly expand in capabilities, the barrier between vast text documents and actionable data will continue to dissipate.

Despite this trend of increased data accessibility, we observed fundamental ’genomic gaps’ where documented chemistry exists from literature without linked sequences. For example, there are 25 BAHD entries related to Phytozome 14 species where the reaction is known but no sequences or accession numbers were extracted from the original papers^6^ (**Fig 1A, Supp. File 2)**. Overall, there are 63 such entries from the FuncFetch-derived data that were not included in FZ-BAHD due to lack of sequence information. Some of these BAHDs may be redundant with known sequences in our database. Even so, their activities are not entirely redundant, 13 unique acceptors are associated with these entries and no other BAHDs **(Supp. File 3)**. When evaluated across substrates, there are 26 unique acceptors and 15 unique donors that have no association with sequenced BAHDs. This gap is concentrated in a few categories of acceptors with more than one of these compounds: Isoflavonoids (5), Phenylpropanoids (C6-C3) (4), Tyrosine alkaloids (4), Flavonoids (3), Diterpenoids (3), and Small peptides (2) (**Fig. 1B**). The lack of sequence-level associations with these substrates presents a barrier for comprehensive functional annotation of this enzyme family. Without this vital connection, these documented activities are inaccessible to sequence-based prediction techniques. We also reiterate a notable caveat that BAHDs whose functions were characterized using only genetic techniques were not included in FZ-BAHD, therefore homologs of proteins such as Glossy2, Glossy2-like, and CER2 involved in processes such as cuticular wax biosynthesis cannot be detected using the database.

FuncPred-OG revealed that the BAHD functional knowledge gap extends beyond missing sequence links. Even when sequences can be assigned to orthologous groups, several OGs lack any characterized representative, preventing direct function transfer by sequence similarity. This is especially notable for deeply conserved but uncharacterized OGs, whose broad conservation suggests biological relevance despite their absence from the experimental record. These OGs are high-value targets for future characterization because validating even one representative could provide functional context for many related enzymes. In this regard, FuncPred-OG serves not only as a prediction tool, but also as a conservation-aware map of where experimental work could most efficiently expand enzyme annotation. This remaining orthology gap further motivates FuncPred-AI, which complements conservation-based inference by proposing substrate-class hypotheses for enzymes lacking characterized relatives.

Nevertheless, FuncPred-AI remains dependent on the same experimental record that limits orthology-based prediction and is therefore constrained by the sparsity, imbalance, and incompleteness of known enzyme activities. To maintain stable model evaluation, we only trained classifiers for labels that contained >5 positive examples across all five cross-validation folds (**Fig. 5D**). As a result, several class labels present in the test set, including Limonoids, Betalain alkaloids, and Yohimbine-type alkaloids, could

not be evaluated by FuncPred-AI. This limitation is compounded by the promiscuous nature of BAHD catalysis. For each binary classifier, enzymes with a documented activity for a given substrate class of interest were treated as positive examples, whereas enzymes without a documented activity were treated as negatives. However, many of these apparent negatives may include possible true-positive enzyme-substrate combinations that have not been tested so far, rather than being confirmed cases of biochemical inactivity. Future work could address this issue using positive-unlabeled learning approaches, which are designed for settings with reliable positives but many ambiguous unlabeled examples, a common challenge in bioinformatics^47^. Alternatively, systematic substrate-screening datasets, such as recent work testing 38 *Arabidopsis* aminotransferases against a shared substrate panel, provide experimentally grounded positive and negative results for the same enzymes and may offer a stronger basis for training substrate-specificity models^48^. Despite these shortcomings, our results show that 93.8% of the FuncPred-AI predictions above a probability score of 0.9 were correct at the top-3 threshold. We note that this high rate was only obtained with 16 enzymes with eight substrate classes, and may change as more training data and more novel enzymes are included.

While FuncPred-AI’s predictive performance is one of the primary outcomes of this work, the underlying protein representations also provide insight into the types of information captured by different PLM feature spaces. ESMC sequence embeddings supported donor and acceptor prediction despite being derived from sequence alone, suggesting that sequence-trained PLMs capture information relevant to enzyme specificity. ESM3 function logits also showed strong separation of OGs and donor types in the unnormalized PCA/Fisher Discriminant Ratio analysis (**Fig. 5A**), although this effect was reduced after normalization and should therefore be interpreted cautiously. The ESM3:FL classifier nevertheless achieved useful performance for acceptor class prediction (**Fig. 5C**), indicating that the function logit representation contains signal relevant to substrate specificity. This is notable because the ESM3 function track was trained using broad InterPro- and GO-derived functional annotations, not experimentally resolved substrate-specific labels or the fine biochemical distinctions that separate closely related enzymes within a family^20^. These results are consistent with recent protein modeling studies showing that textual or semantic descriptions of protein function can improve protein representations and support transfer to downstream tasks. Examples include ProtST^49^ and ProtNote^50^ for text-guided protein function prediction, ProteinDT^51^, ProtDAT^52^, and ProtET^53^ for text-conditioned protein design or editing, and embedding-fusion approaches that combine complementary PLM representations^54^. Together, these studies suggest that semantically informed representations can capture functional information not always available from sequence embeddings alone. Additionally, this information may be unevenly distributed within a model: layer-wise analysis of ESMC shows that functional organization, measured by k-nearest-neighbor accuracy on EC classes, peaks in intermediate layers rather than the final layers that best encode tertiary structure^41^. Because EC-class organization is closely analogous to our substrate-class task, extracting features from intermediate layers may be a promising direction for improved predictive performance and interpretability. Future PLMs trained on denser, experimentally grounded enzyme-function descriptions may therefore improve family-level functional annotation.

Finally, it is useful to contextualize FuncPred within the broader landscape of enzyme function prediction and protein-ligand modeling. At one end of this landscape are mechanistic approaches such as molecular docking, enzyme-focused docking workflows, binding-kinetic modeling, and structural enzymology, which provide mechanistic insight into how a specific enzyme interacts with a substrate. Emerging machine learning approaches address broader-scale enzyme annotation and specificity prediction, including DeepECtransformer for sequence-based EC annotation, EZSpecificity for enzyme-substrate specificity prediction, CLIPZyme for enzyme-reaction alignment in virtual screening, and EnzymeCAGE for structure- and reaction-aware enzyme modeling^19,23,24,55^. FuncPred occupies a complementary position within this landscape. Rather than resolving the detailed physical basis of individual enzyme-substrate interactions or model enzyme-substrate interactions at scale agnostic of family and substrate information, it uses curated biochemical activities and protein representations to map functional potential across a large, specific plant enzyme family. This family-level framework is intended to prioritize candidates derived through experimental assays, assign probable donor and acceptor class specificities, and guide experimental validation by prioritizing high provenance of the experimental data. In predicting substrate classes instead of substrates, FuncPred recognizes the property of enzymes – especially those in specialized metabolism – to promiscuously utilize structurally similar substrates and show allelic/duplicate functional divergence. Once a small subset of substrate classes is identified using FuncPred, the above-mentioned protein-ligand interaction modeling tools can be used to quickly screen a range of relevant substrates from those classes. Together, family-level annotation frameworks, ligand- and reaction-aware ML models, and mechanistic structural approaches can form a closed-loop strategy for enzyme discovery: broad functional prediction nominates candidates, substrate- or reaction-aware models refine hypotheses, and biochemical or structural experiments test and mechanistically explain the predicted activities.

## METHODS

### Curation of BAHD activities

BAHD-family membership was assigned by screening enzyme sequences against the PF02458 profile HMM with hmmsearch using the gathering cutoff (cut –gu). The verified set of BAHD acyltransferases (https://tools.moghelab.org/funczymedb) was manually reviewed for correctness and supplemented with compound metadata. The resulting tabular files were read directly into a set of JSON files (FuncZymeDB) via custom Python scripts. To generate a normalized set of curated compounds, all unique compound names were merged into sets of their canonical name and any number of alternative names based on domain expertise of the authors and compounds structures or identifiers present in papers.

### Compound annotation

Compound annotations were added during construction of the enzyme-compound database by importing a curated compound table and creating one record per compound with names and associated chemical metadata, including PubChem CID, ChEBI ID, InChIKey, and manual donor type (aliphatic/aromatic) annotations. Compound entries with shared curated identifiers or updated annotations were consolidated into single records. Compounds with SMILES were submitted to the NPClassifier API endpoint (https://npclassifier.gnps2.org/classify?smiles=<SMILES>) to append chemical category annotations.

### Network analysis

BAHD donor-acceptor relationships were summarized from the curated enzyme-compound database as unique donor-to-acceptor-Superclass relationships **(Supp. File 1C)**. Acceptor compounds were grouped by NPClassifier Superclass; when a Superclass annotation was unavailable, the NPClassifier Pathway annotation was used instead and denoted with “(p)”. For each donor-Superclass pair, the number of unique acceptor compounds was counted and used to construct a donor-by-acceptor-superclass matrix. For **Fig. 2A**, this matrix was converted to binary presence/absence values and rows and columns were clustered independently using scipy.cluster.hierarchy.linkage with method=“average” and metric=“euclidean”; plotted row and column order was then set with scipy.cluster.hierarchy.leaves_list. Donors were included in the heatmap if they were connected to at least three unique acceptor superclasses. For **Fig. 2B**, donor-superclass relationships were represented as an undirected bipartite graph using networkx. Graph, with donor compounds and acceptor Superclasses or Pathway fallbacks as the two node types. Edges were weighted by the number of unique acceptor compounds supporting each donor-Superclass relationship. Network coordinates were generated with networkx.spring_layout using the edge-weight attribute, method=“energy”, k=1, gravity=1, and a fixed random seed.

### Generation of OGs and inclusion in FuncZymeDB

Following OG generation by OrthoFinder from 85 plant species with an inflation index of 1.5 and other default settings **(Supp. File 4)**, all characterized BAHD enzymes were searched against the uncharacterized sequence set by BLAST, and each characterized enzyme was assigned the OG of its best-matching uncharacterized sequence. These OG assignments were then integrated with species lineage information across 12 manually defined taxonomic levels to generate the OG component of FuncZymeDB **(Supp. File 4)**, which was used for downstream lineage-aware analyses and mapping of characterized and uncharacterized BAHD enzymes (**Fig. 4A-C**).

### Phylogenetic reconstruction

FuncZymeDB BAHD sequences were supplemented with two assayed BAHD enzymes and aligned with MAFFT (v7.520-with-extensions), yielding a final alignment of 338 amino acid sequences^56^. A maximum-likelihood phylogeny was inferred with IQ-TREE 3.0.1 using ModelFinder for substitution model selection and 1,000 ultrafast bootstrap replicates^57^. ModelFinder selected Q.PLANT+R7 as the best-fit model under BIC. The bootstrap consensus tree was visualized in iTOL^58^.

OG annotations were overlaid on the phylogeny by mapping each tree tip to its assigned OG. For each OG represented by two or more tree tips, the least common ancestor was calculated and used to define clade-range annotations in iTOL; single-tip OGs were retained as terminal annotations. Higher-level BAHD clade ranges were generated from a curated OG-to-clade map by calculating the least common ancestor of the OG nodes assigned to each clade. Additional iTOL datasets encoded donor class and acceptor superclass metadata for characterized and assayed enzymes.

### Mapping of characterized BAHDs to OGs using FuncPred-OG

Using a user-selected enzyme family and one or many (1000 maximum) protein amino acid sequences in fasta format, FuncPred-OG then uses BLASTP^16^ (v2.16.0) or mmseqs2^59^ to match each sequence to an OG. In this work, BLASTP was used. If a query sequence was mapped to a characterized OG, another BLAST search was done against all characterized sequences in that OG, and all hits were ranked by percent identity. For each characterized enzyme, donor and acceptor substrates, species, and DOI of paper the activity was reported. These results are displayed on the FuncPred-OG website (https://tools.moghelab.org/funczymedb) and downloadable in a tabular output. The constituent species, narrowest lineage, and mean pairwise identity of the OG are displayed. FuncPred-OG was applied on BAHD sequences from all 85 species, and these results are also downloadable.

### Enzyme cloning and enzyme assays

Full-length coding sequences (CDSs) of *SlSpmHT* and *SlEPS1* were amplified from cDNA of tomato (*Solanum lycopersicum* cv. Ailsa Craig) using primers listed in **Supp. File 6** and cloned into pET28b(+) vector by Gibson assembly. Codon-optimized *ZmPHT* and *OsBEBT* sequences were synthesized and cloned into pET28b(+) by GenScript Biotech Corporation (Piscataway, NJ, USA). All recombinant plasmids were verified by DNA sequencing and transformed into *Escherichia coli* Rosetta 2(DE3) cells for heterologous protein expression. Protein expression was induced by culturing cells at 28 °C with 0.5 mM isopropyl β-D-1-thiogalactopyranoside (IPTG) for 20 h. Cells were harvested by centrifugation at 4000 x g for 5 min at 4 °C and resuspended in 100 mM phosphate buffer supplemented with 1 mM phenylmethylsulfonyl fluoride (PMSF) and 1x protease inhibitor cocktail. The cell suspension was lysed by sonication (24 kHz, 5s on, 30 s off, five cycles), followed by centrifugation at 14000 x g for 15 min at 4 °C. The resulting supernatant was collected as crude enzyme extract and stored at -20 °C until further use.

Enzyme assays were performed with crude enzyme extract in 50 µl reaction mixtures containing 50 mM phosphate buffer (pH 7.2), 50 mM NaCl, 100 µM acceptor substrate and 300 µM donor substrate. The specific combinations of acceptor and donor substrates for each enzyme were provided in **Supp. File 6B**. Crude enzyme extracts prepared from *E.coli* cells carrying empty pET28b(+) vector were included as negative controls to exclude background activity. Reactions were incubated at 30 °C for 1 h and terminated by the addition of 100 µl stop solution consisting of 40% acetonitrile, 40% isopropanol, and 20% water (v/v/v), supplemented with 0.1% (v/v) formic acid, 15 mM telmisartan. After centrifugation at 14000 x g for 10 min, the supernatant was transferred to insert-equipped glass vials and stored at -20 °C prior to liquid chromatography–mass spectrometry (LC-MS) analysis.

Reaction products were detected by LC-MS following previously described procedures^12^. In general, LC-MS analyses were performed using a Dionex Ultimate 3000 HPLC system coupled to a Q Exactive HF Orbitrap mass spectrometer (Thermo Scientific, Waltham, MA, USA). Samples (5 µl) were injected onto an Acquity UPLC HSS C18 column (1.8 mm, 2.1 mm ’ 100 mm; Waters, Milford, MA, USA) maintained at 40 °C. Separation was achieved at a flow rate of 0.4 ml/min using solvent A (water containing 0.1% [v/v] formic acid) and solvent B (acetonitrile containing 0.1% [v/v] formic acid). The gradient program was as follows: 5% solvent B for 1 min, increased linearly to 95% solvent B over 5 min and held for 1min, followed by re-equilibration to 5% solvent B for 0.5 min. Reaction products were detected using parallel reaction monitoring (PRM) methods with predicted precursor ion masses in either positive or negative ionization mode under the following parameters: scan resolution, 17,500; isolation window, 2.0 m/z; collision energies: 20, 40, 60 eV. Data were analyzed using FreeStyle 1.8 SP2 QF1 software (Thermo Scientific). Product peaks were identified based on their exact masses using default peak detection parameters **(Supp. File 6B)**.

### Structure retrieval

Protein structures were collected by first carrying forward structures already linked to characterized sequences in the curated database. Candidate uncharacterized sequences were then deduplicated against the characterized set, and only nonredundant sequences were queried further. UniProt accessions were resolved from existing identifiers, sequence-matched UniProt/UniParc records, or, when needed, by species-constrained BLASTP recovery of GenBank protein accessions followed by UniProt mapping. Structures were then retrieved in priority order from experimentally determined PDB entries linked through UniProt, followed by AlphaFold models (https://alphafold.ebi.ac.uk/api/prediction/<UniProtAccession>) and then SWISS-MODEL repository files (https://swissmodel.expasy.org/repository/uniprot/<UniProtAccession>.pdb). Any missing structures from the characterized set were then generated using AlphaFold version v2.3.2 on Cornell University BioHPC using one A100 GPU^60^. Two of the 336 sequenced characterized BAHDs contained ambiguous amino-acid residues, lacked available structures from the searched resources, and failed AlphaFold structure prediction; these sequences were therefore excluded from structure-dependent feature generation and subsequent FuncPred-AI model training.

### Predicting BAHD substrate categories using FuncPred-AI

ESM-derived feature generation was performed on the subset of characterized BAHDs with complete amino-acid sequences and usable structure files. After exclusion of the two structure-failed ambiguous sequences described above, the model feature cache contained 334 characterized BAHDs, partitioned into 307 training sequences and 27 held-out test sequences. The test set was supplemented with 2 BAHDs assayed in this work for a total of 29 **(Supp. File 7)**. For structure-based ESM3 inputs, each selected PDB file was parsed as a ProteinChain object and converted to an ESMProtein object using ESMProtein.from_protein_chain. For sequence-based inputs, FASTA sequences were loaded directly as ESMProtein objects. ESM3-open (esm3-open) was used to generate structure embeddings and function logits from structure-derived ESMProtein objects, while ESM Cambrian 600M (esmc_600m) was used to generate sequence embeddings from sequence ESMProtein objects.

Raw ESM outputs were first cached as per-enzyme tensors. During model training, cached tensors were loaded and converted to enzyme-level feature matrices before classifier fitting. ESM3 and ESMC embedding tensors were mean-pooled across protein length. ESM3 function-logit tensors were mean-pooled across residues while retaining the function-token axis, and the resulting token-by-feature matrix was flattened into one vector per enzyme. These pooled feature matrices were then used for PCA, standardization, hyperparameter tuning, and classifier evaluation.

Donor models were trained using two non-exclusive binary labels corresponding to aromatic and aliphatic donor type. Acceptor models were trained using NPClassifier-derived acceptor class, superclass, and pathway labels from the curated compound annotations. To improve model stability during hyperparameter tuning and evaluation, acceptor labels were included only if they had at least five positive examples in the training set.

Binary classifiers were implemented with scikit-learn LogisticRegression in a one-versus-rest framework. Feature matrices were standardized with StandardScaler, and class imbalance was handled with class_weight=“balanced”. During cross-validation, standardization parameters were fit on the training set and applied to the test set. For final model bundles, the scaler was fit on the full training matrix and serialized with each classifier so that query sequences are transformed using the same training-set parameters before prediction.

Logistic-regression hyperparameters were selected with GridSearchCV using five-fold cross-validation and micro-averaged AUPR as the refit metric. The search included liblinear models with L1 or L2 penalties and lbfgs models with L2 penalty. Regularization strength C was searched across all orders of magnitude from 1e-4 to 1e4, and tolerance was searched across all orders of magnitude from 1e-6 to 1e0; models were fit with max_iter=5000.

Training-set performance was evaluated by leave-one-out cross-validation. To assess sensitivity to sequence redundancy, additional evaluations were performed after CD-HIT clustering of training sequences at 95%, 90%, 80%, and 70% identity, using singleton sequences at each threshold as the evaluated subset^29^ **(Supp. Fig. 3A,B)**. Performance was summarized using AUPR and AUROC. Final classifiers and their associated standardization objects were serialized as pickled scikit-learn model bundles for downstream FuncPred-AI inference.

## ACKNOWLEDGMENTS

We thank Cornell Bioinformatics Facility and members of the Buckler, Thiede, and Moghe labs at Cornell University and the Busta lab at the University of Minnesota Duluth for helpful advice and feedback during the project.

## AUTHOR CONTRIBUTIONS

NSS and GDM designed the study; NSS led the computational experiments; XY conducted wet-lab experiments; GDM, CM, CG and SS assisted NSS and XY with the curation, data analysis, and website design and implementation; NSS, GM and XY wrote the manuscript; all authors read and approved the final version.

## DATA AVAILABILITY

FuncPred predictions can be downloaded from https://tools.moghelab.org/funczymedb All relevant code has been deposited on GitHub (https://github.com/n8smith-io/FuncZymeDB).

## SUPPLEMENTARY MATERIAL

Supplementary Figure 1: Principal component analysis (PCA) of raw BAHD feature representations.

Supplementary Figure 2: Principal component analysis (PCA) of L2-normalized BAHD feature representations.

Supplementary Figure 3: Effects of sequence identity threshold, acceptor label level, and training set positives on classifier training performance.

Supplementary Figure 4: Pre-correction test-set views of acceptor class confusion and Hit Rate @k.

Supplementary Figure 5: Feature generation resource use for ESM3:FL-SE compared to ESMC:SeqEmbed.

Supplementary Figure 6: Coverage of uncharacterized BAHD predictions by species and FuncPred method.

Supplementary File 1: BAHD activities, compounds, and reactions.

Supplementary File 2: Species-level counts of characterized BAHD enzyme entries in FuncZymeDB, partitioned into unique-sequence, redundant-sequence, and no-sequence records, with Phytozome 14 membership indicated.

Supplementary File 3: Unique acceptors associated exclusively with Phytozome 14 BAHD entries lacking linked sequences, with corresponding enzyme-entry counts, IDs, and species.

Supplementary File 4: A list of species used in this study.

Supplementary File 5: Pairwise sequence-identity and functional-overlap of BAHD clades and Orthologous Groups.

Supplementary File 6: Information on tested enzymes. Supplementary File 7: Information on BAHD test set.

