## Supplemental Fig. 1 for "Systematic functional annotation of thousands of BAHD acyltransferases in plant genomes using Protein Language Model and phylogenomic tools"

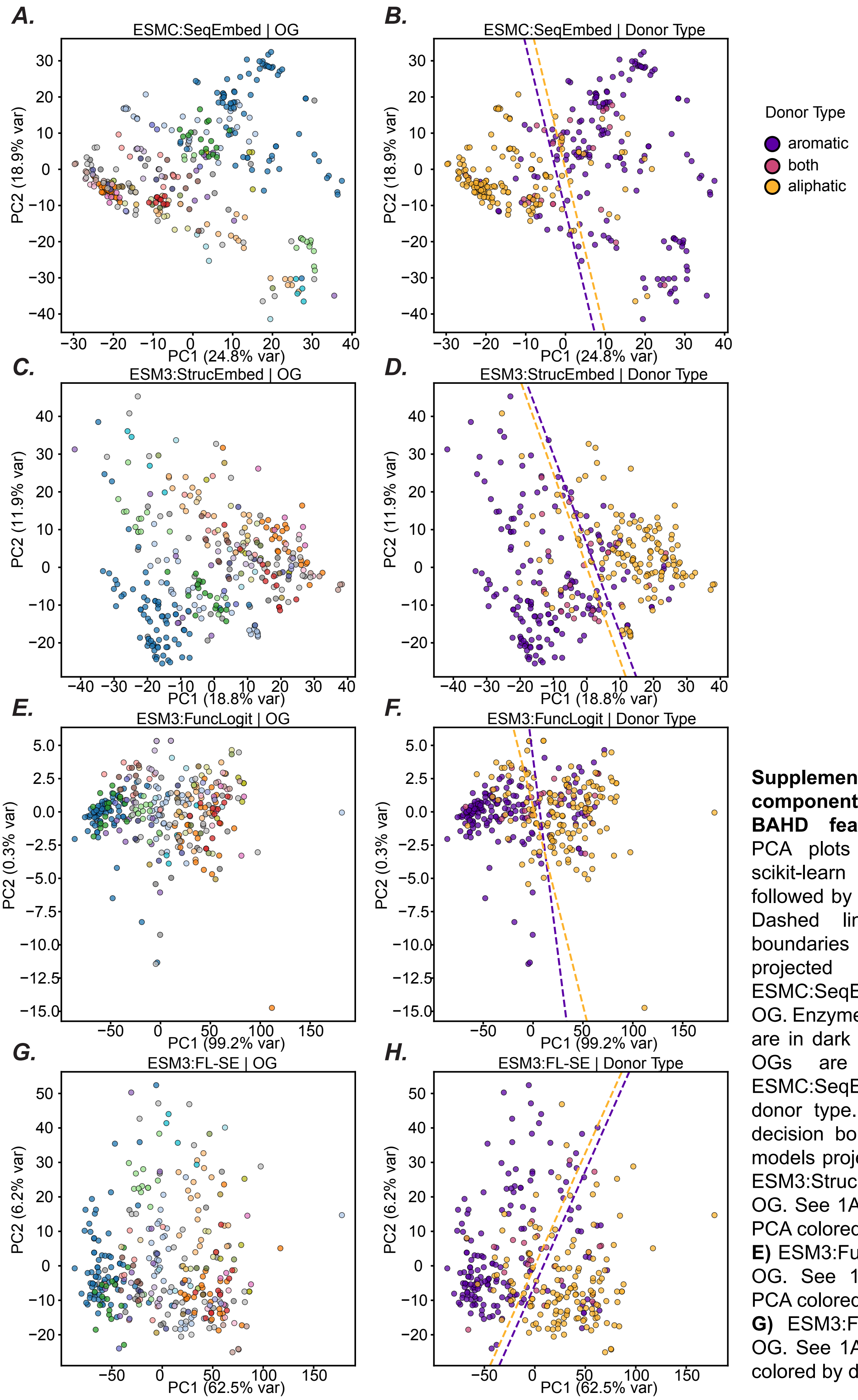


**Supplementary Figure 1: Principal component analysis (PCA) of raw BAHD feature representations.** PCA plots were generated with scikit-learn using StandardScaler followed by PCA on the feature set. Dashed lines indicate decision boundaries of donor type models projected into PC space. **A)** ESMC:SeqEmbed PCA colored by OG. Enzymes without assigned OGs are in dark grey. Singleton enzyme OGs are in light grey. **B)** ESMC:SeqEmbed PCA colored by donor type. Dashed lines indicate decision boundaries of donor type models projected into PC space. **C)** ESM3:StrucEmbed PCA colored by OG. See 1A. **D)** ESM3:StrucEmbed PCA colored by donor type. See 1B. **E)** ESM3:FuncLogit PCA colored by OG. See 1A. **F)** ESM3:FuncLogit PCA colored by donor type. See 1B. **G)** ESM3:FL-SE PCA colored by OG. See 1A. **H)** ESM3:FL-SE PCA colored by donor type. See 1B.
