## Supplemental Fig. 2 for "Systematic functional annotation of thousands of BAHD acyltransferases in plant genomes using Protein Language Model and phylogenomic tools"

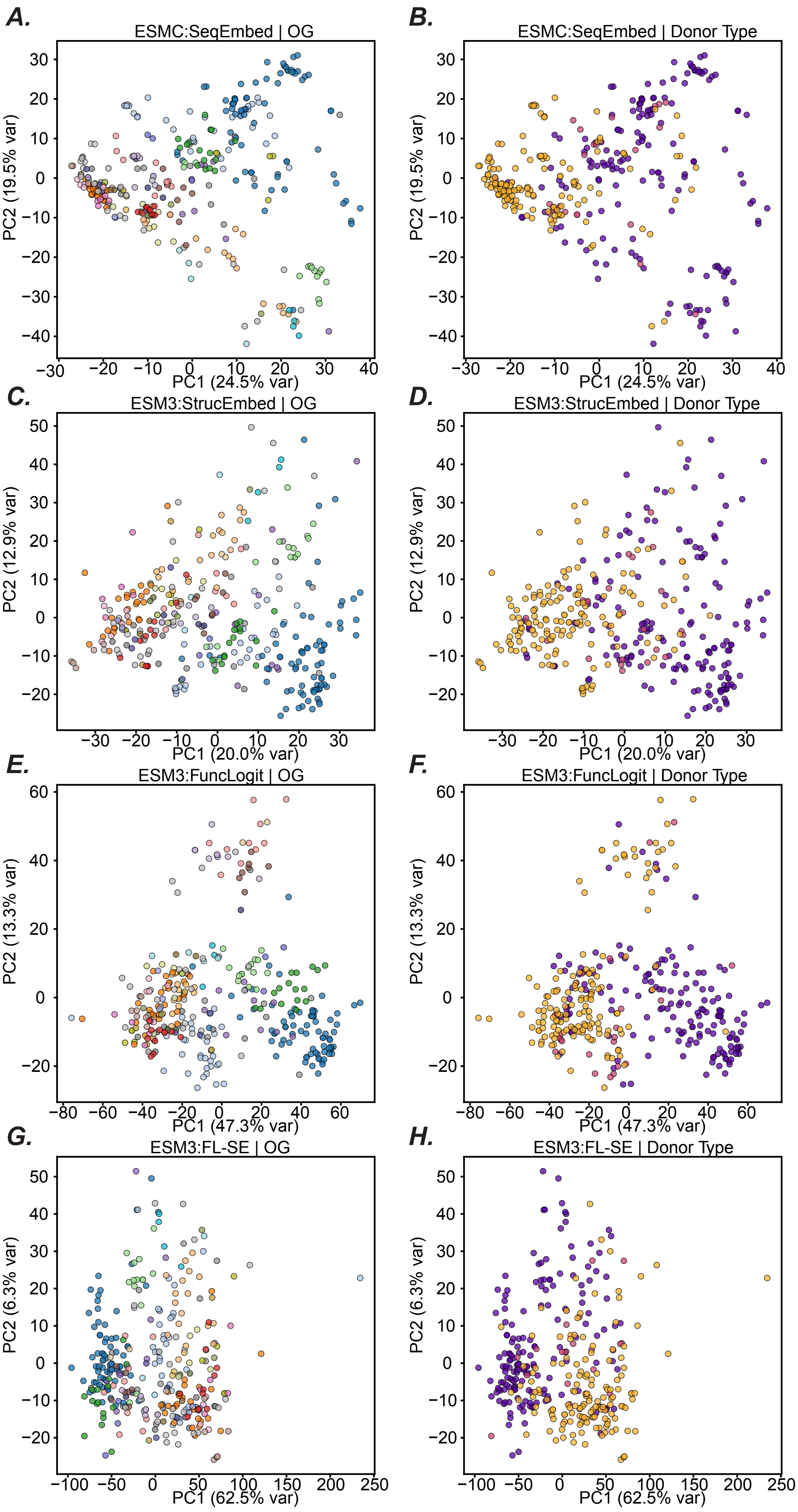

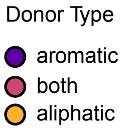


**Supplementary Figure 2: Principal component analysis (PCA) of L2-normalized BAHD feature representations.** PCA plots were generated with scikit-learn using L2 normalization of each feature vector, followed by StandardScaler and PCA on the feature set. Dashed lines indicate decision boundaries of donor type models projected into PC space. **A)** L2-normalized ESMC:SeqEmbed PCA colored by OG. Enzymes without assigned OGs are in dark grey. Singleton enzyme OGs are in light grey. **B)** L2-normalized ESMC:SeqEmbed PCA colored by donor type. **C)** L2-normalized ESM3:StrucEmbed PCA colored by OG. See 2A. **D)** L2-normalized ESM3:StrucEmbed PCA colored by donor type. See 2B. **E)** L2-normalized ESM3:FuncLogit PCA colored by OG. See 2A. **F)** L2-normalized ESM3:FuncLogit PCA colored by donor type. See 2B. **G)** L2-normalized ESM3:FL-SE PCA colored by OG. See 2A. **H)** L2-normalized ESM3:FL-SE PCA colored by donor type. See 2B.
