## Supplemental Fig. 3 for "Systematic functional annotation of thousands of BAHD acyltransferases in plant genomes using Protein Language Model and phylogenomic tools"

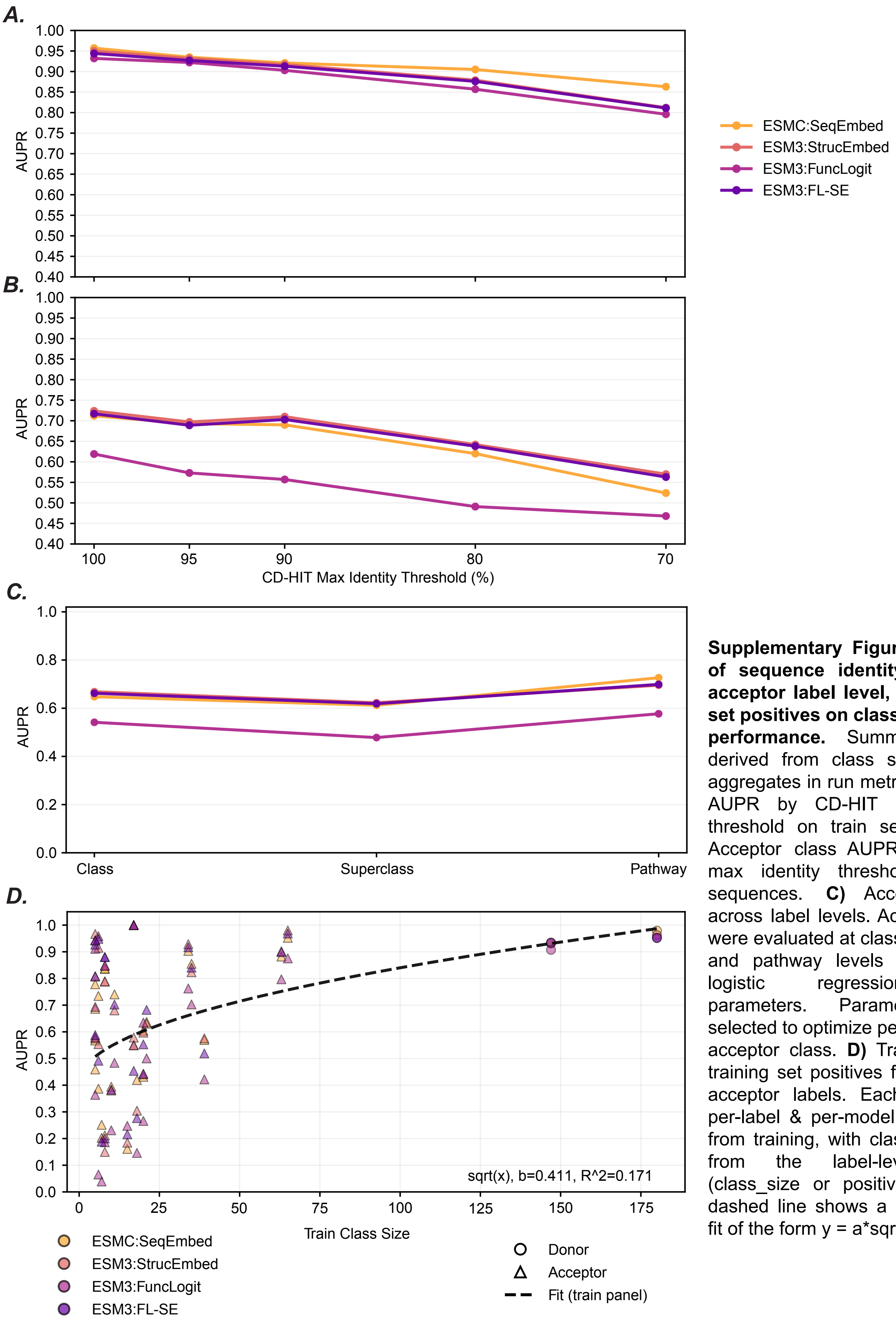


**Supplementary Figure 3: Effects of sequence identity threshold, acceptor label level, and training set positives on classifier training performance.** Summary metrics derived from class size weighted aggregates in run metrics. **A)** Donor AUPR by CD-HIT max identity threshold on train sequences. **B)** Acceptor class AUPR by CD-HIT max identity threshold on train sequences. **C)** Acceptor AUPR across label levels. Acceptor labels were evaluated at class, superclass, and pathway levels using frozen logistic regression model parameters. Parameters were selected to optimize performance on acceptor class. **D)** Train AUPR by training set positives for donor and acceptor labels. Each point is a per-label & per-model AUPR value from training, with class size taken from the label-level metrics (class_size or positive_total). The dashed line shows a least-squares fit of the form y = a*sqrt(x) + b.
