## Supplemental Fig. 4 for "Systematic functional annotation of thousands of BAHD acyltransferases in plant genomes using Protein Language Model and phylogenomic tools"

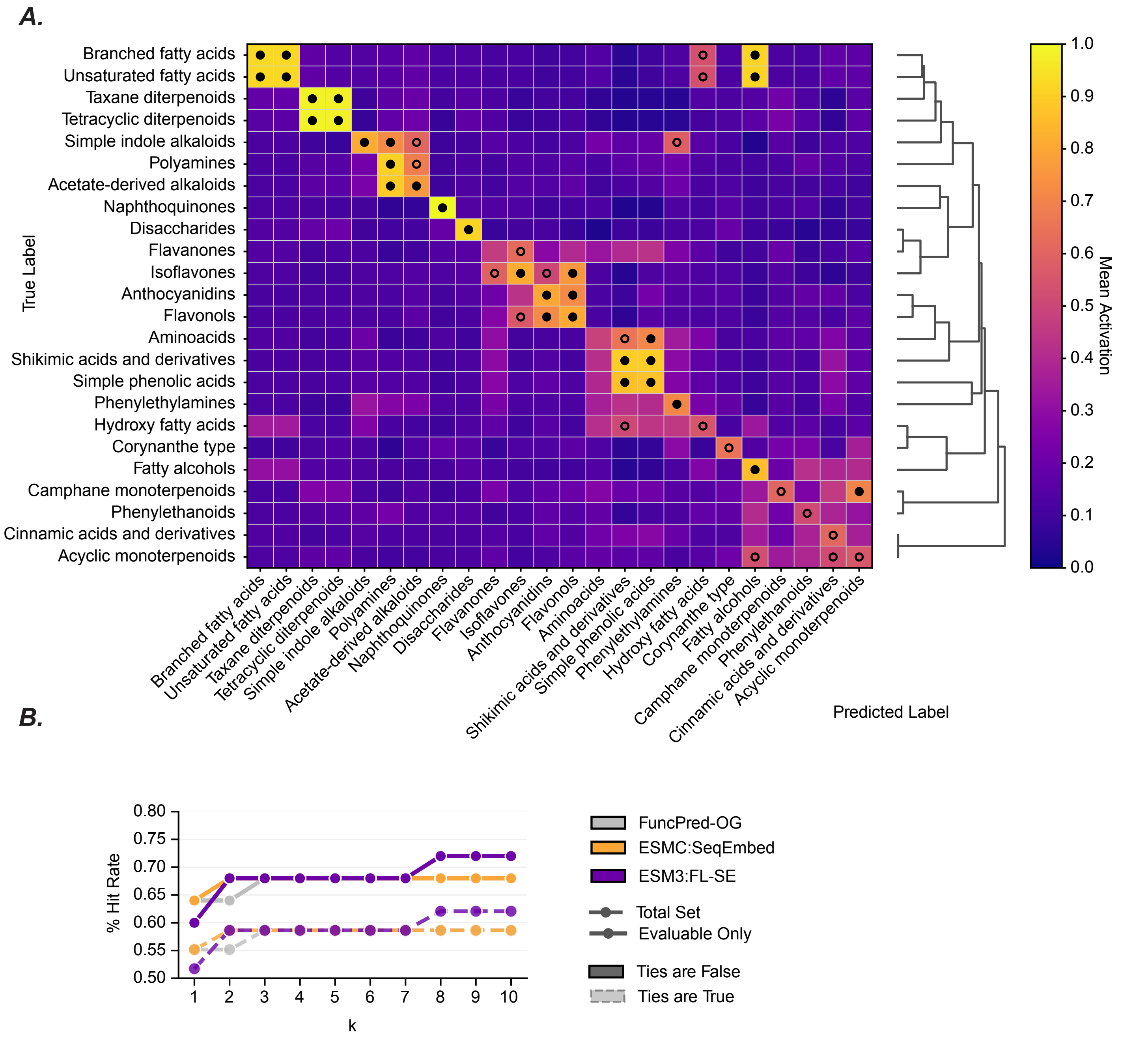


**Supplementary Figure 4:** Pre-correction test-set views of acceptor class confusion and Hit Rate @k. See Main Figure 6A and 6D for the corresponding post-correction results. Main Figures 6B and 6C were unchanged by correction. **A)** Pre-correction test-set cross-label heatmap for ESMC:SeqEmbed acceptor class predictions. Each cell shows the mean prediction activation for the column label among test-set enzymes with the row label. Labels were hierarchically clustered with SciPy using Euclidean distance on row activation profiles after replacing NaN values with 0.0, followed by average-linkage clustering. The same clustered order was applied to both axes. Hollow black circles mark cells with mean activation > 0.5. Filled black circles mark cells with mean activation > 0.7. The post-correction test-set heatmap shown in Main Figure 6A was derived from this panel by merging acceptor classes into connected groups when both off-diagonal directional activations for a label pair were >= 0.7. After grouping, prediction scores for merged labels were collapsed by taking the maximum score within each merged group, and the corrected activation matrix was replotted as Main Figure 6A. **B)** Pre-correction acceptor class prediction Hit Rate @k on the held-out test set. Queries were defined as test-set enzymes. A hit was defined as at least one true label within the top k predicted labels. Ties at the kth boundary were retained. Solid lines indicate the evaluable-only subset, defined as queries for which at least one true label was present in that method's possible outputs, and dashed lines indicate the full common query set. The paired post-correction plot in Main Figure 6D uses the same query set and scoring rules.
