## Supplemental Fig. 5 for "Systematic functional annotation of thousands of BAHD acyltransferases in plant genomes using Protein Language Model and phylogenomic tools"

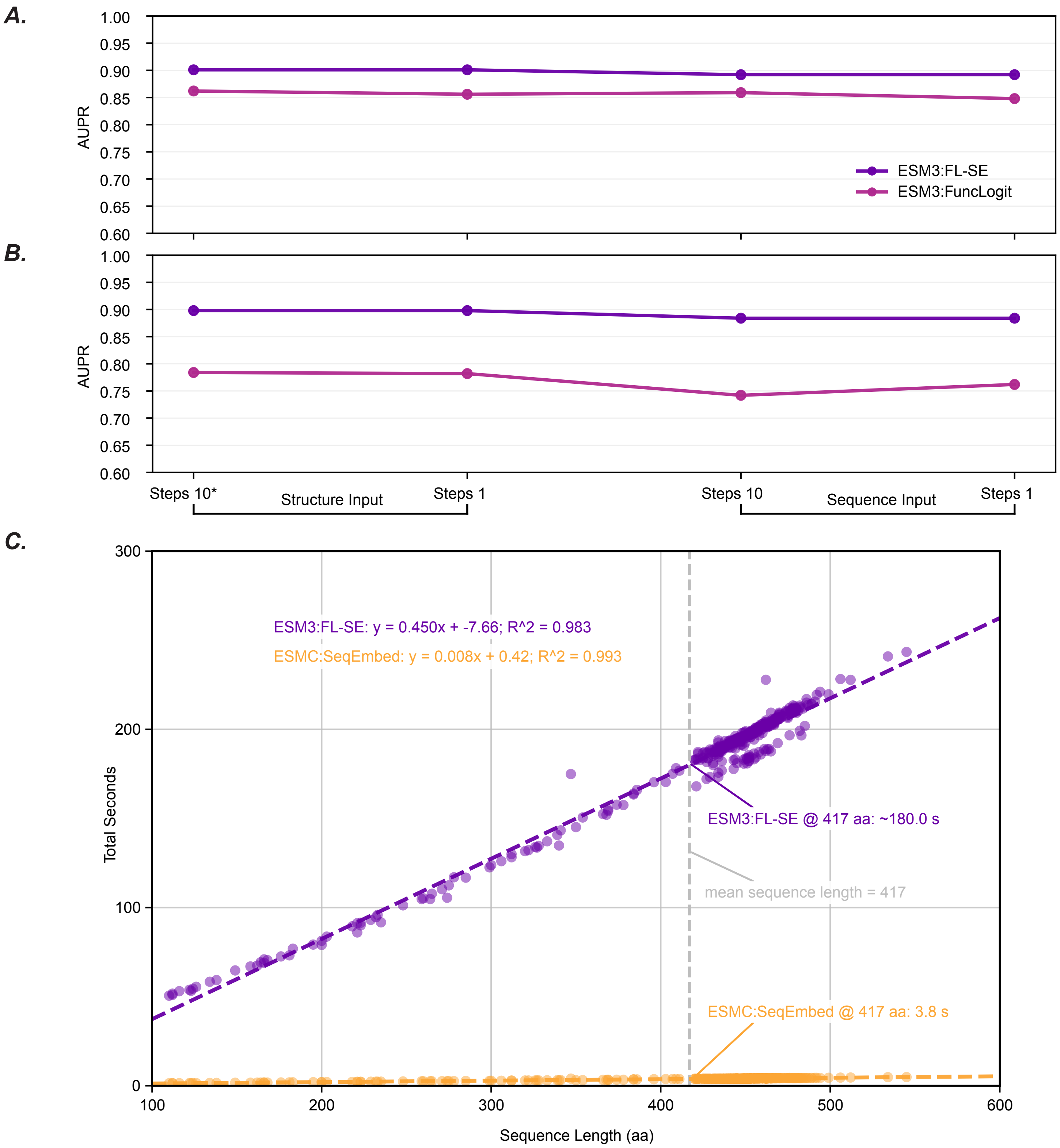


**Supplementary Figure 5: Resource use and performance of ESM3:FL-SE compared to ESMC:SeqEmbed. A)** Donor AUPR on test set by input variant. The asterisk denotes the feature variant used during training. The structure-embedding component differs only by input (structure or sequence), whereas the function-logit component differs by input and the number of ESM3 generation steps (10 or 1). X-axis brackets from 5B apply to this plot. **B)** Acceptor class AUPR on test set by input variant. See 5A. **C)** Total feature generation time by sequence length for selected uncharacterized BAHD sequences. Timing of generation was integrated into a Python script. ESM3:FL-SE time was calculated as the sum of the separately timed ESM3 structure-embedding and ESM3 function-logit generation steps for the same sequence. Both features were generated from the same input structure context, but not as a single joint timed pass. Sequences longer than 600 amino acids were excluded (n=380). Dashed lines show linear least-squares fits of total seconds versus sequence length.
