## Supplemental Fig. 6 for "Systematic functional annotation of thousands of BAHD acyltransferases in plant genomes using Protein Language Model and phylogenomic tools"

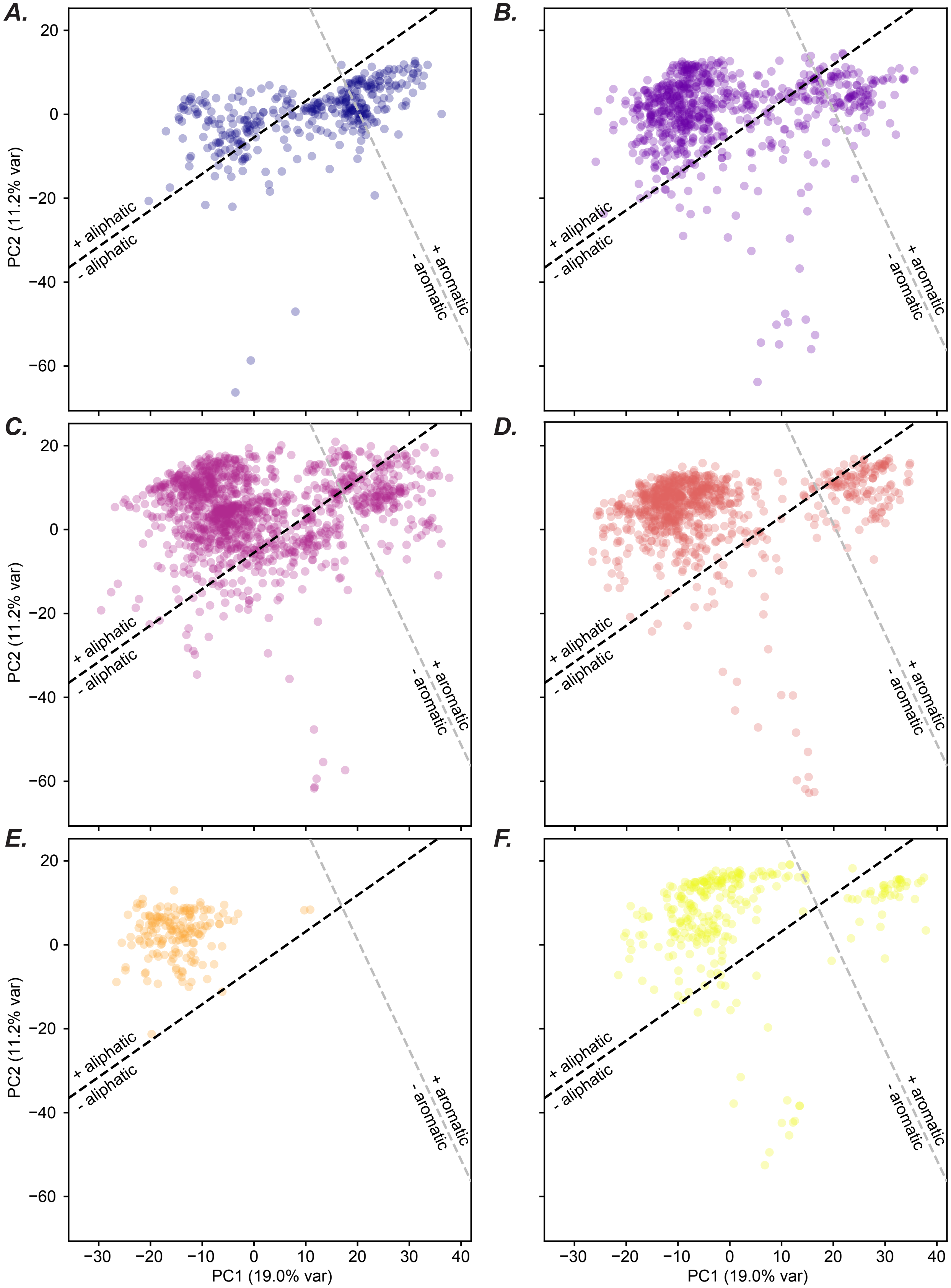


**Supplementary Figure 6: Principal component analysis (PCA) of uncharacterized BAHD ESMC:SeqEmbed features by clade.** Plots generated with scikit-learn using StandardScaler followed by PCA on the uncharacterized ESMC:SeqEmbed feature set. Only uncharacterized sequences assigned to OGs with clade labels were included (See Fig. 4). Dashed lines indicate aromatic and aliphatic donor type model decision boundaries projected into PC space. Clades were ordered and colored based on the fraction of characterized enzymes in each clade that interact with only aromatic donors (e.g. Clade 4 (A, darkest) is most aromatic associated and Clade 7 (F, lightest) least). **A)** ESMC:SeqEmbed PCA of uncharacterized BAHD sequences assigned to Clade 4 (n=354). B) ESMC:SeqEmbed PCA of uncharacterized BAHD sequences assigned to Clade 5 (n=733). C) ESMC:SeqEmbed PCA of uncharacterized BAHD sequences assigned to Clade 6 (n=1316). D) ESMC:SeqEmbed PCA of uncharacterized BAHD sequences assigned to Clade 1 (n=881). E) ESMC:SeqEmbed PCA of uncharacterized BAHD sequences assigned to Clade 3 (n=167). F) ESMC:SeqEmbed PCA of uncharacterized BAHD sequences assigned to Clade 7 (n=282).
