## Supplemental Fig. 7 for "Systematic functional annotation of thousands of BAHD acyltransferases in plant genomes using Protein Language Model and phylogenomic tools"

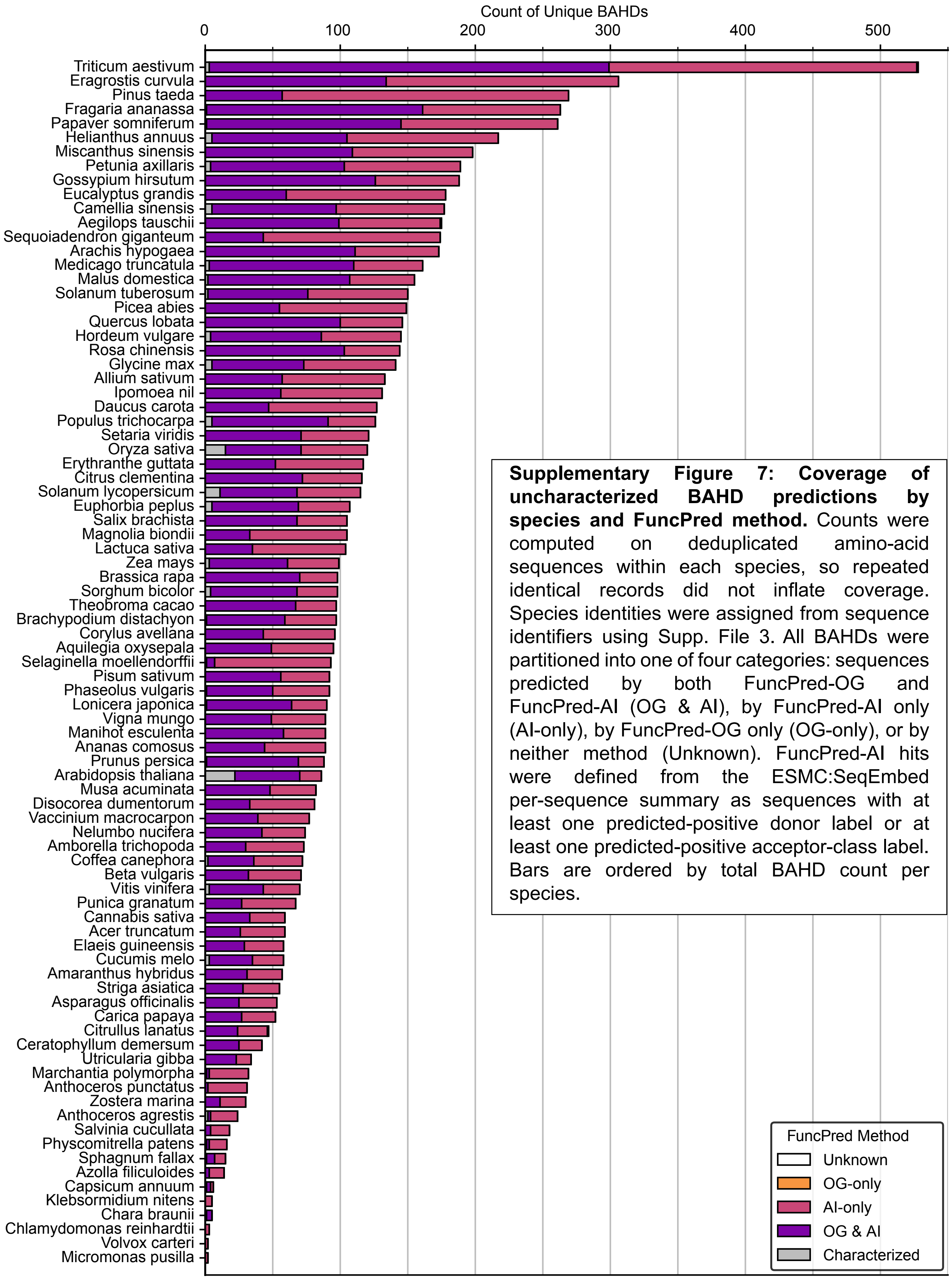


**Supplementary Figure 7: Coverage of uncharacterized BAHD predictions by species and FuncPred method.** Counts were computed on deduplicated amino-acid sequences within each species, so repeated identical records did not inflate coverage. Species identities were assigned from sequence identifiers using Supp. File 3. All BAHDs were partitioned into one of four categories: sequences predicted by both FuncPred-OG and FuncPred-AI (OG & AI), by FuncPred-AI only (AI-only), by FuncPred-OG only (OG-only), or by neither method (Unknown). FuncPred-AI hits were defined from the ESMC:SeqEmbed per-sequence summary as sequences with at least one predicted-positive donor label or at least one predicted-positive acceptor-class label. Bars are ordered by total BAHD count per species.
